# Modeling the simultaneous dynamics of proteins in blood plasma and the cerebrospinal fluid in human *in vivo*

**DOI:** 10.1101/2023.12.28.573559

**Authors:** Pierre Giroux, Jérôme Vialaret, Jana Kindermans, Audrey Gabelle, Luc Bauchet, Christophe Hirtz, Sylvain Lehmann, Jacques Colinge

## Abstract

The analysis of protein dynamics or turnover in patients has the potential to reveal altered protein recycling such as in Alzheimer disease, and to provide informative data regarding drug efficacy, or certain biological processes. The observed protein dynamics in a solid tissue or a fluid is the net result of protein synthesis and degradation, but also transport across biological compartments. We report an accurate 3-biological compartment model able simultaneously account for the protein dynamics observed in blood plasma and the cerebrospinal fluid (CSF) including a hidden central nervous system (CNS) compartment. We successfully applied this model to 69 proteins of a single individual displaying similar or very different dynamics in plasma and CSF. This study put a strong emphasis on the methods and tools needed develop this type of model. We believe it will be useful to any researcher dealing with protein dynamics data modeling.

## INTRODUCTION

Brain research and in particular investigations interested in brain diseases have identified the cerebrospinal fluid (CSF) as a convenient source of information regarding the central nervous system (CNS) proteome (Bastos *et al*., 2017). Indeed, the CNS is in close contact with the CSF through the CNS-CSF barrier, and it exports a large number of proteins to the CSF. Other CSF proteins originate from blood, which is also in close contact with the CSF at the choroid plexus through the blood-CSF barrier. These proteins can be imported by the CNS. That is, alterations of the CNS functioning are likely to yield an alteration of the CSF proteome, which is accessible for diagnosis purposes. Numerous studies hence searched for CSF biomarkers of brain disorders, degenerative diseases predominantly (Bader *et al*., 2020; Karayel *et al*., 2022; Johnson *et al*., 2023). An even more accessible body fluid for patient diagnosis is blood, which naturally led to search for other biomarkers that would go to circulation through the blood-brain barrier directly or through the CSF (Leuzy *et al*., 2022).

Clinical and research proteomics have established powerful methods to determine protein abundance in tissues (Meyer and Schilling, 2017). Parallel and complementary to these efforts, techniques were developed to map the dynamics – or turnover – of proteins (Doherty and Whitfield, 2011). Protein dynamics has the potential to reveal specific disease alterations in protein degradation or clearing. This has been for instance demonstrated for amyloid-β (Aβ), Tau, or sTREM2 in Alzheimer disease (AD) (Mawuenyega *et al*., 2010; Sato *et al*., 2018; Suárez-Calvet *et al*., 2016), retinol-binding protein 4 (RBP4) in diabetes (Jourdan *et al*., 2009). The measure of protein dynamics is commonly performed by mass spectrometry (MS), and it relies on the introduction of an isotopic tracer that labels newly synthesized proteins through a mass shift (Bateman *et al*., 2006; Jaleel *et al*., 2006; Doherty *et al*., 2012; Claydon *et al*., 2012; Wilkinson, 2018). The ratio of labeled versus unlabeled protein MS signals is named the relative isotope abundance (RIA). Protein dynamics parameters are obtained from the change of RIA over time by mathematical modeling.

In this study, we explore how the simultaneous acquisition of proteome dynamics in blood plasma and CSF can be related. In particular, by adapting methods of pharmacokinetics designed to model the diffusion of drugs in the various organs and body compartments, we propose mathematical models including hidden or implicit CNS proteome dynamics. The approach is illustrated using unique unpublished data obtained from a patient with serial blood and CSF collection over 36 hours. Different procedures are available to introduce the isotopic tracer. Here, we applied a protocol that entails the intravenous injection of ^13^C_6_-Leu for nine hours with serial collection of blood and ventricular CSF over an extended period of time (Paterson *et al*., 2019; Lehmann *et al*., 2015), see Figure 1A. It is slightly adapted from the stable isotope labeling kinetics (SILK) protocol (Bateman *et al*., 2006). This pulse-chase protocol both unravels a new protein synthesis phase and its clearance. It reveals patient physiology since protein dynamics in CSF and blood result from potential local synthesis and degradation, but also transport from or to different organs such as the CNS and the liver (Figure 1B).

**Figure 1.**
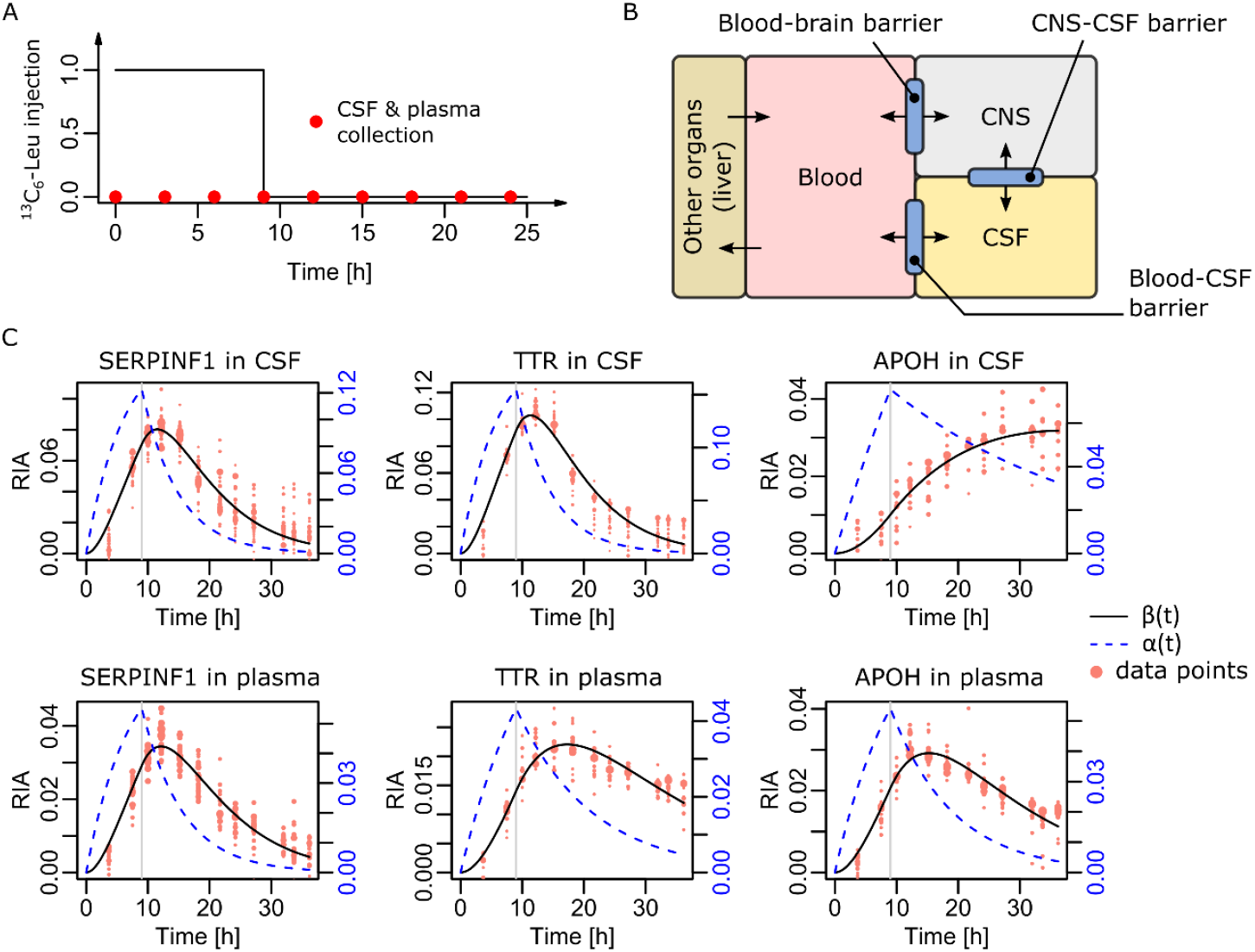
Experimental setting and data available. (**A**) Schematic of the SILK protocol. ^13^C_6_-Leu was injected for the first nine hours, and samples were collected roughly every three hours, starting at *t* = 0 [h]. (**B**) Connections between the considered biological fluids with specific barriers. (**C**) Three typical patterns of simultaneously observed dynamics. Serpin family F member 1 (SERPINF1) harbored comparable dynamics in both CSF and plasma. Transthyretin (TTR) displayed faster dynamics in the CSF. Apolipoprotein H (APOH) displayed the opposite difference with faster plasma dynamics. The black curve *β*(*t*) models the dynamics. The blue dashed curve *α*(*t*) represents the availability of the tracer for the specific protein synthesis. Dot sizes are on an arbitrary scale proportional to the square-root of the ^13^C_6_-Leu-labelled peptide MS intensities. Vertical gray lines at 9 [h] indicate end of tracer injection.

In a previous report, we introduced a mathematical model able to accurately capture protein dynamics in a single tissue at a time (Lehmann *et al*., 2019). The models presented here extend this elementary modeling effort to the multiple biological compartments situation. Interestingly, the respective dynamics in blood and in CSF may display distinct patterns (Figure 1C). This indicates a nontrivial contribution of the CNS to induce the observed dynamics.

## MATERIALS AND METHODS

### Human samples collection

Samples were generated following the clinical protocol “In Vivo Alzheimer Proteomics (PROMARA)” (ClinicalTrials Identifier: NCT02263235), which was authorized by the French ethical committee CPP Sud-Méditerranée IV (#2011-003926-28) and by the ANSM agency (#121457A-11). The enrolled patient (P017) was hospitalized in neurosurgery unit due to subarachnoid hemorrhage (posterior communicating artery aneurysm), and received a temporary ventricular derivation of the CSF. She was 40 years old. Tracer injection and sequential sample collection started 19 days after initial, medical ventricular drainage and normalization of CSF clinical chemistry analysis (protein concentration at 0.35 g/L to compare with normal range 0.2-0.4 g/L range (Roche *et al*., 2008); cell count per mm^3^ was 100). CSF and blood plasma were collected at multiple time points after injection of the tracer (roughly every 3 hours) for 36.2 hours in total. We applied the ethically approved (see above) original SILK ^13^C_6_-Leu infusion protocol (Lehmann *et al*., 2015). Briefly, ^13^C_6_-Leu prepared *per* the European Pharmacopeia (Tall *et al*., 2015) was intravenously administered. After a 10 min initial bolus at 2 mg/kg, an 8h50 infusion at 2 mg kg/h was performed. Ventricular CSF or plasma EDTA samples were collected starting at the beginning of the ^13^C_6_-leucine infusion, roughly every 3h (3 to 6 mL). Samples were transported to the laboratory at 4°C, and centrifuged at 2000g for 10 minutes. CSF and plasma samples was aliquoted into 1.5-mL polypropylene tubes and stored at –80°C until further analysis.

### Sample preparation

1μL of plasma and 150μL of CSF were depleted with depletion columns (High Select™ Depletion Spin Columns, A36370, ThermoFisher). The filtrate was collected and evaporated to dryness on SpeedVac (50 °C). Samples were reconstituted with 20 μL Ammonium Bicarbonate (ABC) 100 mM, 1% SDS and transferred on Eppendorf™ twin.tec™ 96-Well (30129300). Samples were reduced, alkylated and digested with autoSP3 protocol (Müller *et al*., 2020). On AssayMap BRAVO (Agilent), SP3 protocol version 1.0.2 was used. Proteins were reduced with 5μL of Dithiothreitol 80mM during 1800s at 60°C. Then, they were alkylated with 5μL of Iodoacetamide 200mM during 1800s at 30°C. A 50/50 mix of Sera-Mag stock solution A and B was generated at 100 mg/mL and 5μL was added to the sample. 35μL of acetonitrile was added and sample were incubated 1080s. After this incubation, beads were washed two times with 200μL of 80% EtOH and one time with 180μL of acetonitrile. Proteins were digested by adding 35μL of ABC 100 mM, 5μL of Trypsin/LysC (0.05 μg/μL, Promega), and incubate overnight at 37°C and 450 rpm, well closed with sealing foil.

Digestion was stopped with addition of 10μL 5% formic acid. Generated peptides were fractionated on C18 tips (AssayMAP 5 μL Reversed Phase (RP-S) cartridges, G5496-60033, Agilent T) at basic pH. 50μL of 200mM ammonium formate pH10 were added to samples and “Fractionation V2.0” was ran on AssayMap BRAVO (Agilent T). Briefly, cartridges were primed with 100μL of 90% acetonitrile, equilibrated with 50μL 20mM ammonium formate pH10, before sample loading. Cartridges were washed with 50μL 20mM ammonium formate pH10 before sequential elution with 35μL. For the CSF, 5 fractions were generated: 15%, 20%, 25%, 30% and 80% acetonitrile in 200mM ammonium formate pH10. For the plasma samples, 4 fractions were generated: 15%, 20%, 25% and 80% acetonitrile in 200mM ammonium formate pH10. In this condition, the fractions at 15% and 80% were mixed. Fractions were diluted with 0.1% formic acid and loaded on evotip following the manufacturer procedure.

### Chromatography and MS analysis

LC-MS acquisitions were performed on Evosep One using 8cm x 150μm, 1.5μm (EV1109, Evosep) with 60SPD method coupled to TIMS TOF HT (Bruker Daltonics) through a captive spray ion source. Ion source parameters were 1500V on capillary with 3.0L/min at 180°C for the drying gas. DDA-PASEF method was used in positive ion mode. MS1 range was 100-1700 m/z. TIMS settings were 1/K00.75-1.25, Ramp and Accumulation time of 100ms. At MS2 level, 10 PASEF Ramps were performed per cycle of 1.17s. Plasma fractions were analyzed in duplicate.

Data acquisitions were submitted and interrogated inside the Paser Box (Bruker Daltonics). Uniprot database (2021) was used with human as the only taxonomy. Contaminants were added during the database indexation on Paser Box server. CID mode was selected for the fragmentation with monoisotopic precision at precursor and fragment level. Mass tolerance was 20ppm at the precursor level and 30ppm at the fragment level. Precursor mass range was between 600 and 6000 Da. Proteins were digested with trypsin with strict specificity and maximum 2 missed cleavages. Minimal peptide length was set to 6 amino-acids with a maximum of 2 potential variable modifications as deamidation (N, Q) and oxidation (M). Carbamidomethylation was used as fixed modification of cysteine. XCorr was used as primary score, Zscore in the secondary score, and TIMScore was used. A minimum of 1 peptide identified per protein was required. False discovery rate (FDR) less than 1% was imposed at the protein level.

Identification results were exported with mzIdent files (mzid and mgf files). The se files were used to import Peptide Search into skyline. Peptide and precursor ions in the library were uploaded on skyline file. Retention time tolerance was 1.5 min on MS/MS scan ID, 0.2 on ion mobility value coming from the experimental library, 3 isotopes at resolution 60000 at MS1 level. Isotope modification was added for ^13^C_6_-leucine.

### Individual fluid mathematical model

Computing the ratio of tracer-containing MS signals versus the total (tracer-containing and non-tracer-containing) signals defined the RIAs such as illustrated in Figure 1C (salmon dots). Detailed derivation of our 2-compartment mathematical model to fit data from an individual sample was published (Lehmann *et al*., 2019). For clarity, we provide a brief summary. For a given peptide and time point, the observed RIA is defined by the ratio of the heavy Leu signal *H* (observed at +6 Da *per* Leu) and the total signal *L + H, L* being the signal at the nominal mass. The curve traced by RIA values over time is modeled by *β*(*t*). Our 2-compartment model comprises a first compartment representing the rate of tracer availability denoted *α*(*t*). The second compartment represents the rate *β*(*t*) of newly synthesized peptides. Modeling at the protein level is achieved by pooling all the peptide RIA values at all available times, and fitting the same mathematical model on the pooled data. The system of ODEs defining *α*(*t*) and *β*(*t*) is

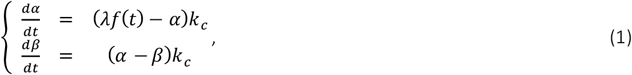

with *α*(0) = 0 = *β*(0). Figure 1C illustrates three typical proteins in each fluid. It is important to observe that the parameter λ essentially acts as a scale parameter, whereas *k*_*c*_, the clearance rate primarily acts as a shape parameter that strongly conditions protein half-life.

A peculiarity in modeling RIAs, which results from noise and the large differences of intensities between *L* and *H* (ratio 10 to 100 usually), is that observed RIA values may contain a slight vertical shift (see SI for an explanation of this phenomenon and Figures S1-S2). Therefore, parameter fitting in (1) must include the computation of a shift *s* along with λ and *k*_*c*_ to adjust *β*(*t*) to the data. Classically, minimizing the summed squared errors between *β*(*t*) and the observed RIAs minus the shift *s* provides the solution. We empirically found that weighing squared errors proportionally to 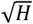lead to better fit since RIAs with stronger *H* signals were more accurate. Minimization was achieved by a quasi-Newton iteration (function optim in R with method BFGS). The system (1) was numerically integrated by RADAU5 method (Hairer and Wanner, 1996) available from R deSolve package (function radau).

Robustness against outlier RIA values caused by noisy data was obtained thanks to a 2-step empirical procedure. A first model was fitted using all the available RIAs, and then RIAs located at a distance larger than half the difference between the minimum and maximum values of the first fitted *β*(*t*) model were considered outliers. A second application of the quasi-Newton method without the outliers was then performed to determine the final parameter values. In addition, RIAs at time 0 were always considered outliers since no tracer incorporation had occurred yet. The bootstrap was used to estimate confidence intervals around parameter estimates. To use RIAs obtained at *t* = 0 to estimate the shift *s* does not work, most likely due to variable co-eluting material.

### Initial data processing pipeline

Plasma and CSF MS data were processed separately, essentially following the method we already published to extract usable spectra and protein dynamics models in each fluid (Lehmann *et al*., 2019). The only differences with respect to this original method was to add additional peptide-level qualitative filters. Since each detected peptide could be present in more than one protein fraction and at different charge states, we name observation a given peptide in a given fraction at a given charge state. Many peptides obviously gave rise to multiple observations. Dynamics estimation is based on observations. Starting with Skyline export, a first step implemented in a Perl script eliminated peptides devoid of leucine. Otherwise, the analysis was conducted in R. Second step was to eliminate observations for which there were too many missing time points or insufficient signal intensity. In a third step, the remaining observations data were fitted with mathematical model (1). The fitted model enabled us to eliminate aberrant shapes, incompatible with protein dynamics and isotopic tracer incorporation. For observations whose shapes were potentially acceptable, we defined a band around the model curve *β*(*t*) to call outliers RIAs that fell outside this band.

Observations harboring too many outliers were eliminated. In addition, we required Spearman correlation ≥ 0.75 between non-outlier data points and *β*(*t*) values at corresponding times. We also used a piecewise polynomial model (loess) to estimate a 95% confidence area around the non-outlier data points, and we required that *β*(*t*) remained within this area at least 75% of the time. Lastly, we imposed that at least two non-outlier data points were available before 10 hours and after 20 hours to constrain initial and final dynamics. The final stage was to combine all the observations available for a protein into a single model. For this purpose, we only considered unique peptides to avoid contamination between proteins. As soon as three or more observations were available, we checked the parameters *k*_*c*_ obtained for each one independently at the previous stage. Observations from unique peptides harboring an outlier *k*_*c*_ value (R function boxplot.stats) were discarded. The remaining observations were aligned on the observation with highest median heavy Leu signals because of the vertical shift issue. Finally, the mathematical model was fit as above on the pooled observations to obtain the protein model complemented by a bootstrap (1,000 times) to estimate parameter 95% confidence intervals (CI95). The steps of this pipeline are detailed in SI.

The pipeline above performed the separate analysis of each fluid. For each fluid, its output consisted in a set of protein models with their parameters and, most importantly, all the non-outlier RIAs of all the corresponding validated observations. These RIAs constitute the input data for the simultaneous CSF-plasma models that are the object of this study as soon as a protein was detected in both fluids.

## RESULTS AND DISCUSSION

### Mass spectrometry data processing

Overall, CSF MS data covered 3,156 proteins and 22,842 distinct peptides, 16,913 of which contained at least one leucine. Each peptide was subjected to filters for signal quality and the requirement of being detected at least 9 out of the 13 time points in the same chromatographic fraction and at the same charge state. This resulted in 2,417 distinct peptides usable for dynamics modeling that corresponded to 3,179 distinct observations, *i*.*e*., a different peptide, chromatography fraction, or charge state. Based on the usable observations, we could determine the dynamics of 869 proteins in the CSF. In plasma, the same process led to 1,260 proteins covered by 9,243 peptides among which 6,788 contained at least one leucine. We found 1,264 usable peptides from 1,740 observations, and obtained an estimation of the dynamics of 271 proteins in plasma. The number of proteins detected with dynamics data in both plasma and CSF was 194.

In this report, with one patient available only, we focused on the ability to model the dynamics of proteins in the CSF and plasma simultaneously. We hence reasoned that we would limit our considerations to the proteins that were available with a rather large number of observations. We imposed a minimum of four observations in each fluid separately, and it reduced the list of common proteins between CSF and plasma from 194 down to 69 (Figures S3-S5).

### Initial considerations

The CSF is rather poor in cells. Most CSF proteins are indeed produced in the CNS, or remote organs such as the liver and brought via the blood. The CSF proteome composition as well as its dynamics are thus defined by the rates of imports and exports through the CNS-CSF and blood-CSF barriers. They marginally depend on CSF local protein synthesis and degradation. Furthermore, remote organs and CNS protein dynamics cannot be measured *in vivo* directly in patients. This forces us to study CSF and plasma protein dynamics with models where CNS and remote organ contributions can only be implicit.

Pharmacokinetics literature describes how to model a compound reaching different body compartments (Bourne, 2018), and a 2-biological compartment model is often applied (Figure 2A). Assuming classical exponential elimination dynamics in each body compartment *C*, written as *C*(*t*) = *Be*^−*bt*^, the model in Figure 2A is represented by the ODE system

**Figure 2.**
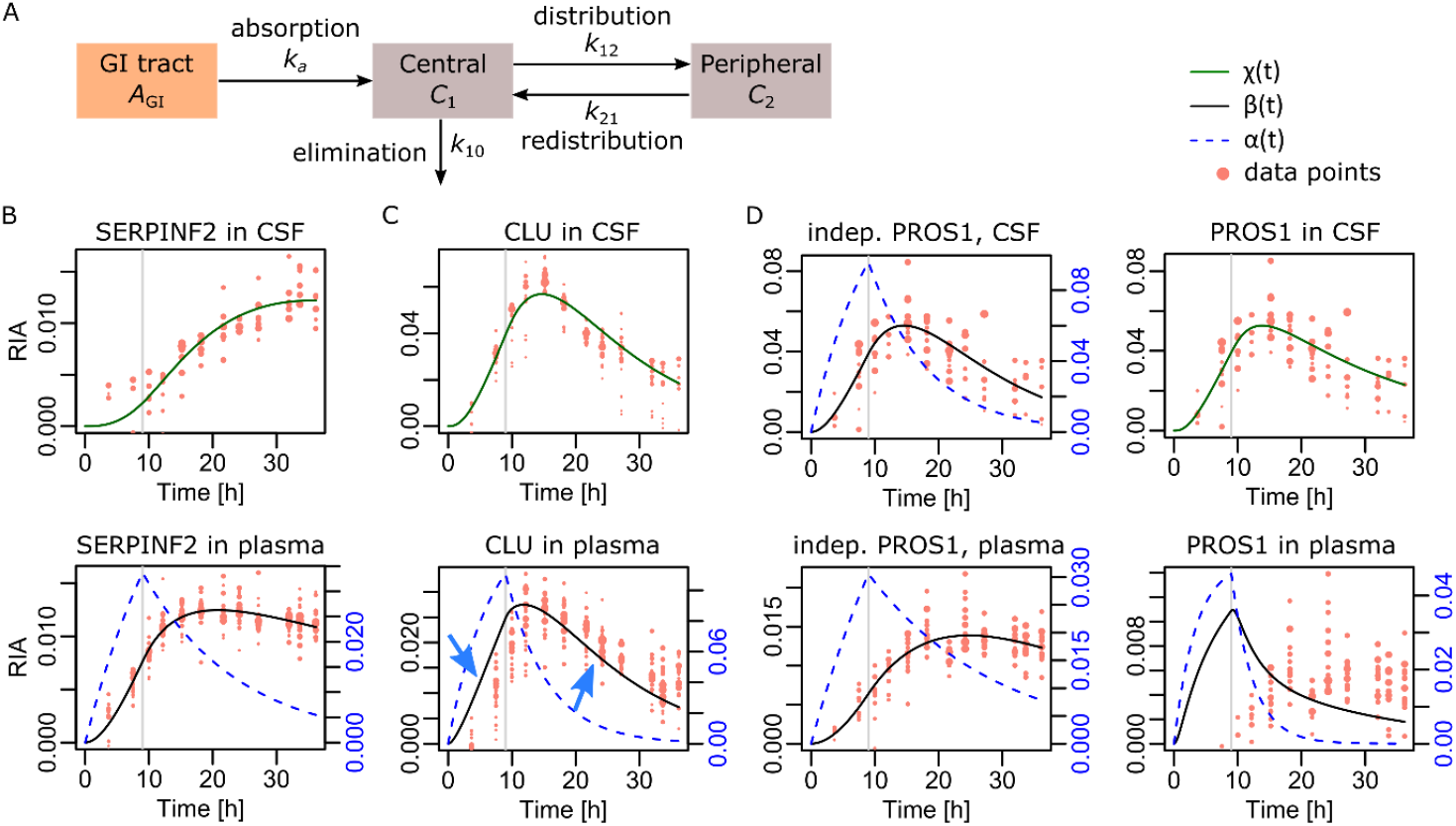
Attempts with a 2-biological compartment model. (**A**) General principle of a 2-biological compartment model in pharmacokinetics. (**B**) Application of such a model to simultaneously capture blood plasma and CSF protein dynamics of Serpin family F member 2 (SERPINF2). (**C**) Application of the model to Clusteri n (CLU). We note the limited accuracy achieved (blue arrows). (**D**) Application of the model to Protein S (PROS1). Left, independent models computed in each fluid separately. Right, the best 2-biological compartment model result we could achieve.

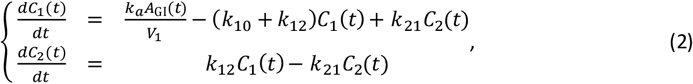

where *C*_1_(*t*) and *C*_2_(*t*) stand for each compartment compound abundance over time, *A*_GI_(*t*) stands for the compound abundance in the gastrointestinal (GI) tract and *V*_1_ for the volume of distribution. This model relates to the 2-compartment model we used to derive (1), but in the latter case we only had *α*(*t*), the infusion of ^13^C_6_-Leu, which is equivalent to *A*_GI_(*t*), and the RIA *β*(*t*), which is equivalent to *C*_1_(*t*). The reason is that there is one additional compartment *C*_2_(*t*) in (2) because pharmacokinetics nomenclature refers to a 2-compartment model as a model with two biological compartments plus the GI tract. Accordingly, in (2), we replace *A*_GI_(*t*)*/V*_1_ by plasma *α*(*t*) and *C*_1_(*t*) by plasma *β*(*t*), and *C*_2_(*t*) by CSF RIA in, which we denote as *χ*(*t*). We hence obtain the system

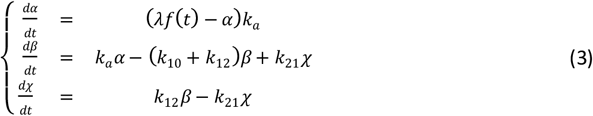

that simply establishes linear transfers between plasma and CSF RIA values at rates *k*_12_ and *k*_21_. For clarity, the system (3) is referred to as a 2-biological compartment model to distinguish from the 2-compartment model (1).

The search for optimal parameters in (3) with an unconstraint quasi-Newton iteration (R optim function, BFGS method) led to non-feasible negative values for some transfer rates. Applying bound-constraint optimization (R optim with L-BFGS-B method) solved this issue. Note that optimization included the vertical shifts on RIAs mentioned in Materials and Methods, one independent shift in each biological fluid. While (3) provided an accurate model for proteins displaying slower dynamics in CSF such as Serpin family F member 2 (Figures 2B and S6A), it sometimes failed for proteins with comparable dynamics in both fluids, for instance Clusterin (Figure 2C). Additional successful examples are featured in Figure S6B. For proteins harboring faster dynamics in CSF, the model (3) systemically failed as for Protein S (Figure 2D) despite trying many initial values for the quasi-Newton iteration. Additional failed examples are featured in Figure S6C.

Before introducing a more complex model, we mention that some authors introduced a notion of delay between biological compartments when modeling protein dynamics (Wildsmith *et al*., 2012). Although this sounds plausible, inspection of all the CSF *β*(*t*) curves in Figures S3-S5 did not reveal any obvious delay. The synthesis of new proteins apparently started immediately within the limits of the accuracy provided by sample collection every three hours. To nonetheless evaluate the potential relevance of introducing a delay, we implemented a modification of (3), where *χ* must instead satisfy

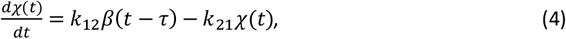

with *τ* > 0, the delay, and *β*(*t*) = 0 for *t* ≤ 0. Eq. (4) inserted in (3) defined a delay differential equation, which we numerically integrated with the function dede of R deSolve package (method set to radau). Parameter search was conducted applying bound-constraint optimization as above. Introducing a delay did not improve the model accuracy, and it caused some instability: slightly different initial values resulted in distinct estimations of *τ* and inaccurate solutions. See Figure S7 for illustrative examples with CLU. Instability was likely induced by excessive parametrization, but since the model was inaccurate there was no point investigating this further.

Overall, the initial considerations above demonstrate that CSF dynamics cannot be generally explained by simply importing proteins from plasma, which was expected and makes a lot of sense physiologically.

### Three-biological compartment models

To be closer to physiology, we introduced an additional biological compartment to the model (3) representing CNS protein synthesis and transport. Namely, we have *C*_1_ = plasma with proteins produces by blood cells and all the organs but the CNS (major protein producer is the liver), *C*_2_ = CNS, and *C*_3_ = CSF (Figure 3A). Following the same mathematical logic that led to system (3), we obtain the *full model*

**Figure 3.**
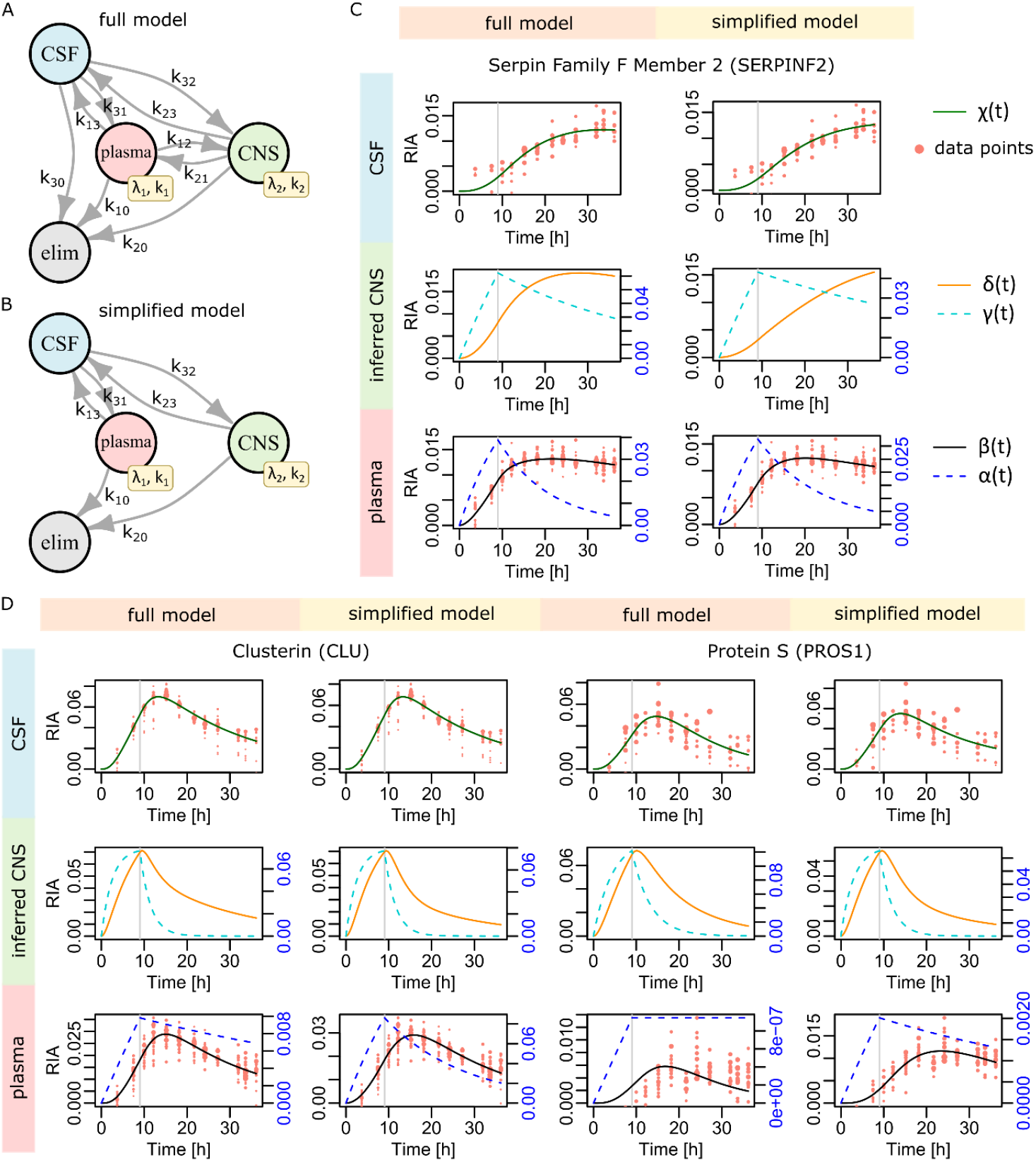
Three-biological compartment models. (**A**,**B**) Model graphical representations. (**C**) Serpin Fami ly F Member 2 (SERPINF2) results. (**D**) Clusterin (CLU) and Protein S (PROS1) results .

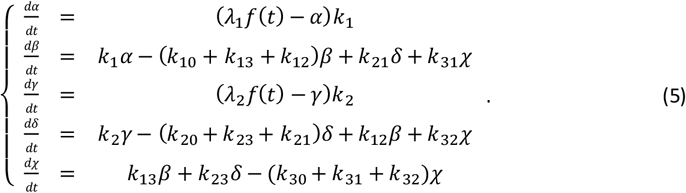

The CNS compartment *C*_2_ is hidden (no experimental data available). In *C*_2_, the functions γ(*t*) and *δ*(*t*) play the same role as *α*(*t*) and *β*(*t*) in *C*_1_. *β*(*t*) must fit plasma data and *χ*(*t*) must fit CSF data. The role and definition of the model parameters is obvious from (3). In case a few proteins would also be synthesizes/degraded in CSF directly, this contribution would be absorbed by *C*_2_ thanks to the linear nature of the model.

The system (5) contains 13 parameters. The absence of direct observations in the CSF might limit our ability to estimate their values or, at least, to obtain reasonably accurate estimates. Moreover, CSF *in situ* degradation rate *k*_30_ is physiologically questionable, though some authors reported its existence (Ranganathan *et al*., 2006; You *et al*., 2005). Mathematically, the parameter *k*_30_ is redundant with transfers from CSF towards plasma and CNS, *i*.*e*., rates *k*_31_ and *k*_32_, which is likely to lead to numerical difficulties. We thus defined a *simplified model* (10 parameters), where we considered the exports from CNS to plasma as already integrated in plasma RIA data, the contribution of plasma proteins to CSF via first entry in the CNS as negligible, and no CSF *in situ* degradation leading to (Figure 3B)

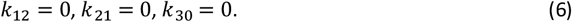

For both models, parameters were searched by bound-constraint optimization as above, including a shift *s*_1_ applied to plasma RIAs, and *s*_3_ to CSF RIAs.

The application of the above two models (full and simplified) to SERPINF2 (Figure 3C) reproduced the accurate fit observed for the independent application of model (3) (Figure 2B). The application of the 3-biological compartment models to CLU (Figure 3D) solved the slight lack of accuracy from which model (3) suffered in Figure 2C. Two more such examples are featured in Figure S8A. Regarding the much more difficult case of PROS1, which dynamics model (3) was unable to capture, we found that the full model (5) experienced similar difficulties. On the contrary, the simplified model (5-6) achieved a perfect fit (Figure 3D). In Figure S8B, we show that both the full and the simplified models accurately fitted data for the other two proteins with faster CSF dynamics that caused difficulties to model (3) in Figure S6C.

These results indicate that the 3-biological compartment approach managed to deliver a general solution to the simultaneous modeling of plasma and CSF protein dynamics. In most cases, the full and simplified 3-biological compartment models yielded accurate solutions. In some cases, the full model led to incorrect solutions, *e*.*g*., for PROS1 (Figure 3D), while the simplified model (5-6) remained accurate. This was most likely due to the removed redundancy between parameters *k*_30_, *k*_31_, and *k*_32_, which otherwise might make parameter fitting of the full model (5) an ill-posed problem (we confirm this in the next section). Accordingly, we decided to keep the simplified model (5-6) as a *bona fide* solution to the simultaneous modeling of protein plasma and CSF dynamics.

### A Bayesian formulation

In the examples above, we used a quasi-Newton iteration to fit the model parameters. This strategy might suffer from a dependency to the initial values used to start the iteration, and it does not provide any information about parameter variability. Although the second issue could be addressed by the bootstrap as we did before (Lehmann *et al*., 2019), initial value dependency would remain unaddressed. We hence decided to apply Bayesian modeling instead, which provides an efficient and natural solution to both issues.

Denoting RIA_*i*_ the *i*^th^ observation in plasma and *β*_*i*_ the corresponding model value, we assume normal errors

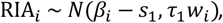

with *i ∈* {1; ⋯ ; *n*}, *n* the number of plasma RIAs, *β*_*i*_ = *β*(*t*_*i*_), *t*_*i*_ the time at which RIA_*i*_ was observed, *w*_*i*_ the weight proportional to 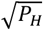for observation *i*, and *s*_1_ the vertical shift of RIAs in plasma. *N*(*μ, τ*) denotes a normal distribution with mean *μ* and precision *τ* (=1/variance). Employing the same notations for RIA_*j*_ and *χ*_*i*_ in CSF, *j ∈* {1; ⋯ ; *m*}, then

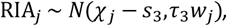

with *s*_3_ and *τ*_3_ the shift and precision in CSF respectively. Further assuming normal priors for the shifts and vague Gamma priors for the precisions, we have (*d ∈* {1; 3})

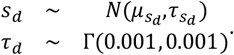

Regarding the many rate parameters, there log-transformed values are modeled with normal priors

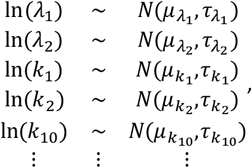

where we set the mean values empirically, or based on the parameters obtained from model (1) applied in each fluid separately. Namely, 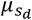was set to 0, ln(λ_1_) mean was set to ln(λ) in plasma, ln(λ_2_) mean to ln(λ) in CSF, ln(*k*_1_) and ln(*k*_10_) means to ln(*k*_*c*_) in plasma, ln(*k*_2_) and ln(*k*_20_) means to ln(*k*_*c*_) in CSF, ln(*k*_13_), and ln(*k*_31_)means to ln(0.1), and ln(*k*_23_) and ln(*k*_32_) means to ln(0.05). Precision 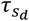was set to 5000, and all the other precisions were set to 10.

The model was coded using BUGS. MCMC parameter sampling was performed with OpenBUGS (Lunn *et al*., 2000), the BUGS code is provided in SI. We found that 200,000 iterations including 100,000 burn-in were sufficient for OpenBUGS safe convergence. We systematically used two Markov chains, and convergence diagnostics was achieved comparing within- and between-chain variability (Brooks and Gelman, 1998). Each chain was initialized with parameter random values drawn from their respective prior distributions, but for the shifts *s*_1_ and *s*_3_ that were initialized with their respective, independent fluid quasi-Newton estimates according to model (1).

In Figure 4A, we report the result of the simplified model with Bayesian parameter estimation for SERPINF2. For this protein, the 2-biological compartment model was already able to capture the simultaneous plasma and CSF dynamics with good accuracy (Figure 2B). This suggested that CNS contribution should remain modest or be associated with rather high parameter variability (no strong constraint). Indeed, Figure 4A shows broad uncertainty on the CNS dynamics (left) and CNS transfer rates (right). That is, in the absence of direct CNS measurement, the model could not exclude CNS contribution, but its precise nature logically remained elusive. In Figure 4B, CLU displays a very different behavior. We know from previous attempt that the 3-biological compartment model was necessary to achieve accurate modeling (Figures 2C, 3D). This translated into well-constrained CNS dynamics (right), and less variable CNS transfer rates (left graphic representation). The higher CNS RIA values and elimination rate *k*_20_ compared to SERPINF2 were also in agreement with a more important role of the CNS in CLU CSF dynamics. Figure 4 further illustrates two examples harboring faster CSF dynamics, PROS1 and transthyretin (TTR). In both cases (Figures 4C-D), we again found constraint CNS dynamics and less variable CNS transfer rates. TTR was more pronounced in this mode, which can be explained by slightly more accurate experimental data, a higher ratio between maximal RIA values in CSF and plasma, and an even faster CSF dynamics compared to PROS1.

**Figure 4.**
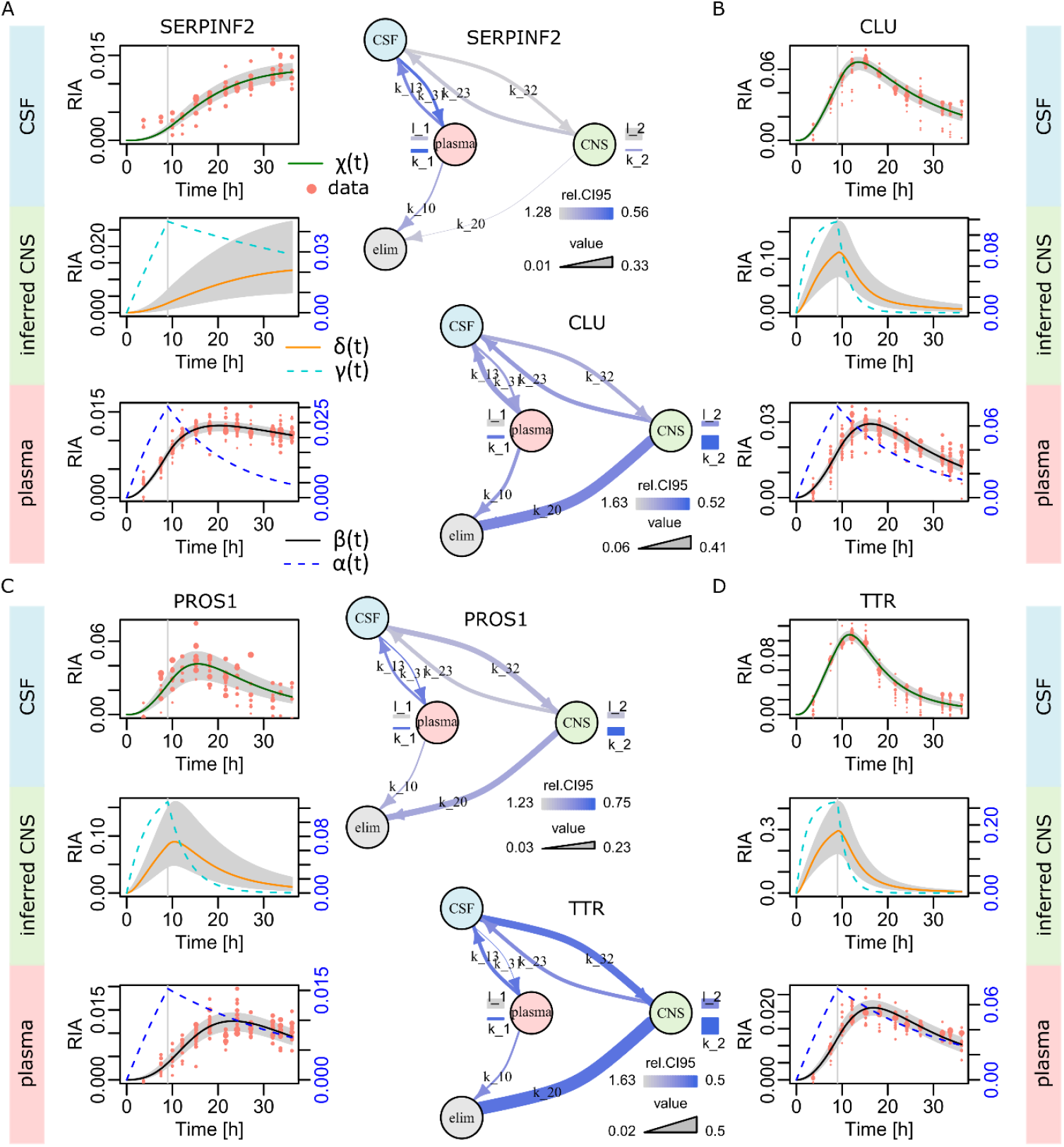
Bayesian inference with the simplified model. (**A**) SERPINF2. Left, the dynamics in the three biological compartments. The gray areas feature the Bayesian estimates of the 95% credibility intervals around *β*(*t*), *δ*(*t*), and *χ*(*t*). Right, graphic representation of the model and its parameters (in the linear space). Parameter magnitude is represented by the line width. Parameter variability is depicted using a color-scale that is based on the relative 95% credibility interval (re.CI95 in the figure), which is the 95% credibility interval range divided by the parameter estimate. (**B**) CLU. (**C**) PROS1. (**D**) Transthyretin (TTR).

To finish this section on Bayesian modeling, we wanted to clarify the reasons for the full model difficulties. In Figure 5A, PROS1 full model is featured and the estimated parameters (solid lines) obviously failed to fit data. The medians of all the *β*(*t*), *δ*(*t*), and *χ*(*t*) curves (dashed lines) generated were much closer to the correct solution. This indicates the existence of multiple solutions in the parameter space that led to equally accurate curves. As a matter of fact, in Figure 5B, plotting the density of the explored (*k*_30_,*k*_31_)- and (*k*_30_,*k*_32_)-spaces, we observe a multimodal distribution. The mean values that were used for parameter estimation (black crosses in Figure 5B) were not aligned with any local maxima in Figure 5B, thereby explaining why Bayesian parameter estimation led to wrong curves. In Figure 5C, we see that similar difficulty happened with the easier SERPINF2 data indicating that the issue is intrinsic to the full model (due to its redundant parameters). In these computations, we set ln(*k*_12_), ln(*k*_21_) and ln(*k*_30_)means to ln(0.1). Increasing the number of iterations from 200,000 to 500,000 resulted in the very same results.

**Figure 5.**
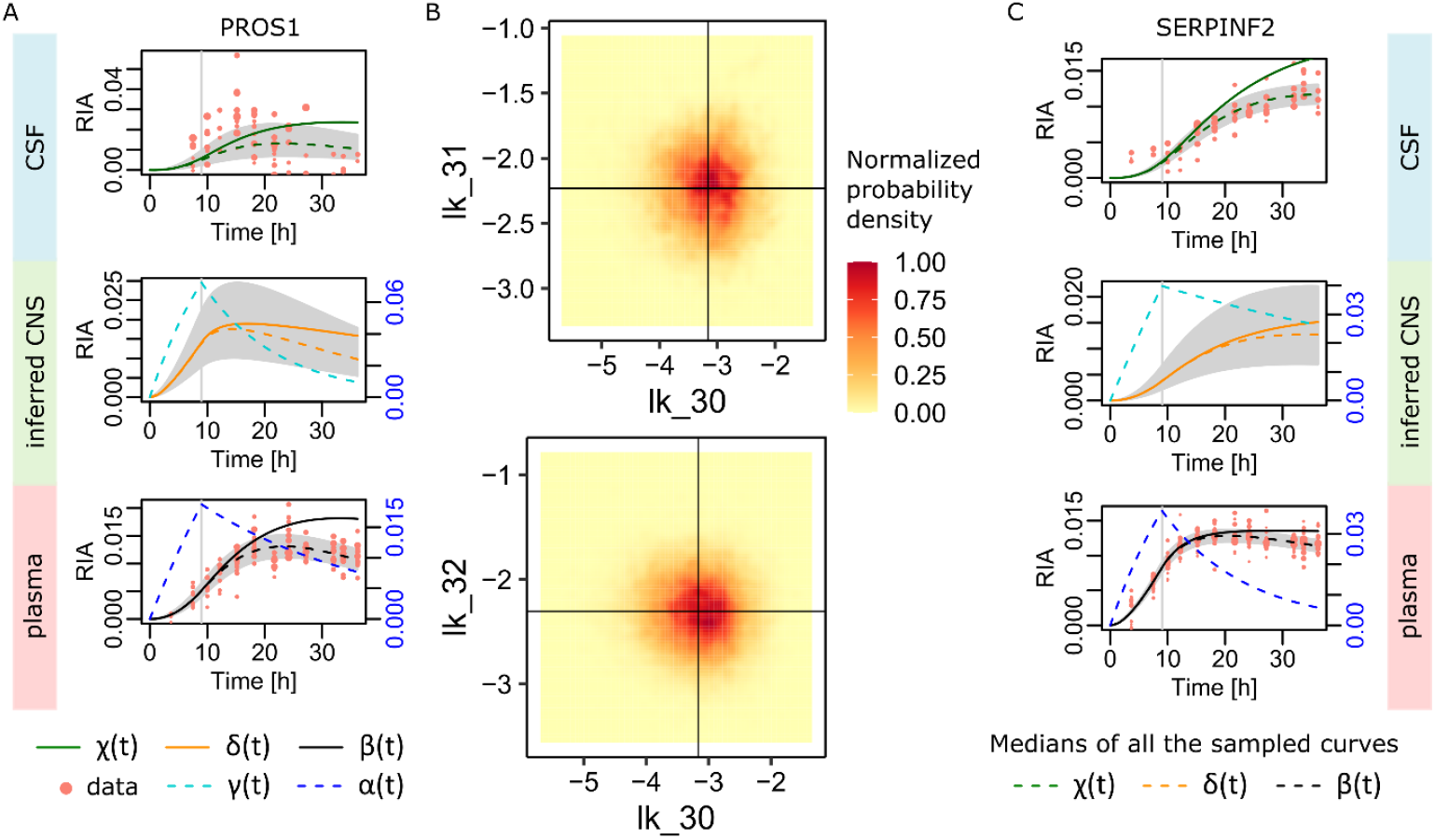
Dissecting the full model difficulties. (**A**) PROS1 dynamics in CSF and plasma were not fit by the full model (5) when applied with the Bayesian estimation of the parameters (soli d lines). The medians of all the curves generated during Gibbs sampling (dashed lines) were correct in plasma and better in CSF. (**B**) The parameter space reduced to the likely redundant transfer rates *k*_30_, *k*_31_, and *k*_32_ . We see a multimodal probability density compatible with the existence of multiple solutions to the parameter fitting problem. Averages used for the solid lines in panel (A) are represented by the black crosses. (**C**) Similar phenomenon on SERPINF2 easier data. In that case, the median curves provided a correct solution, while the solid curves based on the means of the sampled parameters failed to fit data.

## CONCLUSIONS

We have shown that classical pharmacokinetics methodology can be adapted to the problem of modeling protein dynamics in multiple biological compartments simultaneously. This involved first order systems of ODEs with linear transfer rates between biological compartments. The dataset we analyzed was comprised of experimental measures in ventricular CSF and plasma obtained from a human patient *in vivo*. A first result was that although satisfying in some cases, a 2-biological compartment model was not sufficient to account for the observed dynamics of all the detected proteins. CSF physiology makes this fluid a compartment at the interface of blood circul ation and the CNS, but for obvious reasons there was no possibility to acquire protein dynamics data from the CNS directly. Accordingly, a 3-biological compartment model was considered, with the CNS as third (hidden) compartment. Our second main result was that this type of model harbored the necessary flexibility to account for all the observed protein dynamics.

Among the 3-biological compartment models, we considered two variants: a full model with all possible transfers between biological compartments, and a simplified model without transfers between plasma and CNS, and no *in situ* CSF protein degradation. Although one could argue that the full model was physiologically more correct, the estimation of its parameters turned out to be ill - conditioned (multiple solutions due to redundant parameters in the absence of direct CNS measures). Moreover, CSF *in situ* degradation is often considered as marginal despite some reports indicating it might happen in some circumstances (Ranganathan *et al*., 2006; You *et al*., 2005).

Furthermore, transfers between plasma and the CNS that were removed from the simplified model can be regarded as already integrated in the observed plasma data. That is, the simplified model we proposed displayed excellent numerical properties for parameters estimation, it was accurate, and it remains physiologically reasonable. This model combined with Bayesian parameter estimation could precisely capture the dynamics of all the 69 proteins we their different dynamics. The estimated transfer rates between the CSF and CNS compartment reflected the necessity to involve an additional source to plasma when modeling the CSF dynamics.

This very methodological work should provide clear concepts, techniques, and tools for other researchers interested in the dynamics of proteomes and physiology.

## SUPPLEMENTARY DATA

The models for all the 69 common plasma-CSF proteins with at least 4 observations are provided as Supplementary Data along with convergence test results (Brooks and Gelman, 1998), parameters and 95% credibility intervals estimates, and control plots (curves and graphic model representations).

## ACKNOWLEDGEMENTS

JC, SL, and CH were supported by the ANR-20-CE44-0007-01 grant. SL and CH were supported by the PHRC 2010 PROMARA. JC, SL, and CH were supported by the Fondation Alzheimer.

## REFERENCES

Bader, J.M. et al. (2020) Proteome profiling in cerebrospinal fluid reveals novel biomarkers of Alzheimer’s disease. Mol Syst Biol, 16, e9356.

Bastos, P. et al. (2017) Insights into the human brain proteome: Disclosing the biological meaning of protein networks in cerebrospinal fluid. Crit Rev Clin Lab Sci, 54, 185–204.

Bateman, R.J. et al. (2006) Human amyloid-beta synthesis and clearance rates as measured in cerebrospinal fluid in vivo. Nat. Med., 12, 856–861.

Bourne, D.W.A. (2018) Mathematical Modeling of Pharmacokinetic Data 1er édition. Routledge.

Brooks, S.P. and Gelman, A. (1998) General Methods for Monitoring Convergence of Iterative Simulations. Journal of Computational and Graphical Statistics, 7, 434–455.

Claydon, A.J. et al. (2012) Protein turnover: measurement of proteome dynamics by whole animal metabolic labelling with stable isotope labelled amino acids. Proteomics, 12, 1194–1206.

Doherty, M.K. et al. (2012) A proteomics strategy for determining the synthesis and degradation rates of individual proteins in fish. J Proteomics, 75, 4471–4477.

Doherty, M.K. and Whitfield, P.D. (2011) Proteomics moves from expression to turnover: update and future perspective. Expert Rev Proteomics, 8, 325–334.

Hairer, E. and Wanner, G. (1996) Solving Ordinary Differential Equations II: Stiff and Differential-Algebraic Problems 2nd ed. Springer-Verlag, Berlin Heidelberg.

Jaleel, A. et al. (2006) In vivo measurement of synthesis rate of multiple plasma proteins in humans. Am. J. Physiol. Endocrinol. Metab., 291, E190–197.

Johnson, E.C.B. et al. (2023) Cerebrospinal fluid proteomics define the natural history of autosomal dominant Alzheimer’s disease. Nat Med, 29, 1979–1988.

Jourdan, M. et al. (2009) Impact of type 1 diabetes and insulin treatment on plasma levels and fractional synthesis rate of retinol-binding protein 4. J. Clin. Endocrinol. Metab., 94, 5125–5130.

Karayel, O. et al. (2022) Proteome profiling of cerebrospinal fluid reveals biomarker candidates for Parkinson’s disease. Cell Rep Med, 3, 100661.

Lehmann, S. et al. (2019) In vivo large scale mapping of protein turnover in the human cerebrospinal fluid. Anal. Chem.

Lehmann, S. et al. (2015) Stable Isotope Labeling by Amino acid in Vivo (SILAV): a new method to explore protein metabolism. Rapid Commun Mass Spectrom, 29, 1917–1925.

Leuzy, A. et al. (2022) Blood-based biomarkers for Alzheimer’s disease. EMBO Mol Med, 14, e14408.

Lunn, D.J. et al. (2000) WinBUGS - A Bayesian modelling framework: Concepts, structure, and extensibility. Statistics and Computing, 10, 325–337.

Mawuenyega, K.G. et al. (2010) Decreased clearance of CNS beta-amyloid in Alzheimer’s disease. Science, 330, 1774.

Meyer, J.G. and Schilling, B. (2017) Clinical applications of quantitative proteomics using targeted and untargeted data-independent acquisition techniques. Expert Rev Proteomics, 14, 419–429.

Müller, T. et al. (2020) Automated sample preparation with SP3 for low-input clinical proteomics. Mol Syst Biol, 16, e9111.

Paterson, R.W. et al. (2019) SILK studies - capturing the turnover of proteins linked to neurodegenerative diseases. Nat Rev Neurol, 15, 419–427.

Ranganathan, S. et al. (2006) Assessment of Protein Stability in Cerebrospinal Fluid Using Surface-Enhanced Laser Desorption/Ionization Time-of-Flight Mass Spectrometry Protein Profiling. Clin Proteomics, 2, 91–101.

Roche, S. et al. (2008) Clinical proteomics of the cerebrospinal fluid: Towards the discovery of new biomarkers. Proteomics Clin Appl, 2, 428–436.

Sato, C. et al. (2018) Tau Kinetics in Neurons and the Human Central Nervous System. Neuron, 98, 861–864.

Suárez-Calvet, M. et al. (2016) sTREM2 cerebrospinal fluid levels are a potential biomarker for microglia activity in early-stage Alzheimer’s disease and associate with neuronal injury markers. EMBO Mol Med, 8, 466–476.

Tall, M.L. et al. (2015) [Injectable preparation of labeled leucine with the carbon 13 for a clinical research program on the Alzheimer disease: pharmaceutical control of raw materials and the finished product and stability study]. Ann Pharm Fr, 73, 43–59.

Wildsmith, K.R. et al. (2012) In vivo human apolipoprotein E isoform fractional turnover rates in the CNS. PLoS ONE, 7, e38013.

Wilkinson, D.J. (2018) Historical and contemporary stable isotope tracer approaches to studying mammalian protein metabolism. Mass Spectrom Rev, 37, 57–80.

You, J.-S. et al. (2005) The impact of blood contamination on the proteome of cerebrospinal fluid. Proteomics, 5, 290–296.

