## Supplementary Information for "Modeling the simultaneous dynamics of proteins in blood plasma and the cerebrospinal fluid in human *in vivo*"

### Supplementary Methods

#### Origin of vertical shifts

RIA is defined as the heavy leucine-labeled peptide MS signal intensity $H$ divided by the total (light $L$ and heavy $H$) MS signal intensity for this peptide: $\mathrm{RIA} = H/(L + H)$. Now, in the presence of noise, which we supposed to be both additive and multiplicative in full generality, we have an observed ratio given by

$$R = \frac{(1 + r)H + n}{(1 + r)(L + H) + 2n}$$

with $r \geq 0$ the rate of multiplicative noise and $n \geq 0$ the additive noise level. The light peptide signal $L$ is intense compared to $n$. The heavy peptide signal $H$ can be small compared to $L$ and thus commensurate with $n$ in the worst cases. Developing the observed ratio, we obtain

$$\frac{(1 + r)H + n}{(1 + r)(L + H) + 2n} = \frac{H}{(L + H) + \frac{2n}{1 + r}} + \frac{n}{(1 + r)(L + H) + 2n}$$

$$= \frac{H\left(L + H - \frac{2n}{1 + r}\right)}{(L + H)^2 - \left(\frac{2n}{1 + r}\right)^2} + \frac{n}{(1 + r)(L + H) + 2n}$$

$$\cong \frac{H}{L + H} + \frac{-2Hn}{(L + H)^2} + \frac{n}{(1 + r)(L + H) + 2n}$$

Because $2n/(1 + r) \cong 2n$ remains very small compared to $L$ and thus to $L + H$. We now have

$$R \cong \mathrm{RIA} - \frac{2n \times \mathrm{RIA}}{(L + H)} + \frac{n}{L + H} = \mathrm{RIA} + \varepsilon$$

with $\varepsilon = \frac{n}{L+H} - \frac{2n \times \mathrm{RIA}}{(L+H)}$ dependent on the data, *i.e.*, on the peptide abundance and leucine labeling rate mainly, but small and rather constant since $H$ is typically 10 to 100 times smaller than $L$. That is, a small – almost constant – shift on the RIA values is expected, see Figure S1 for simulations with realistic parameters.

An example of peptide RIAs without correction for their individual shifts is featured in Figure S2 below for Alpha-2-Macroglobulin (A2M). See step 4 of the pipeline below for how the individual shifts were removed to pool peptide data at the protein level.

**Initial data processing pipeline**

Step 1: Eliminate peptide devoid of leucine

Step 2: Technical filtering

Initial filtering first regrouped the MS peaks from the same spectra, i.e. the light and heavy signal of a same peptide in a same LC-MS spectrum, as Skyline exports individual peaks on separate lines. Then, the most intense isotope was identified according to the majority rule: most frequently the most intense isotope over times 10.0 to 36.17 in the light species. The maximum intensity of the selected isotope was subsequently required to be higher than $10^4$ (light species again). This threshold was empirically determined to ensure reasonably strong heavy peptide signals, i.e. above noise. Finally, we imposed to have at least 9 out of 13 time points with a minimum of one pair of light/heavy signals available.

Step 3: Construction of a peptide (or observation) model

For each observation that passed step 2, we used ODE (1) in the main paper to fit the observation RIAs. This used the robust procedure described in the main paper Materials and Methods. We imposed the following criteria to accept the shape of the resulting model:

a) $\alpha(t)$ and $\beta(t)$ increasing at $t = 3.67$ ;
b) $\alpha(t)$ decreasing at $t = 12.17$ ;
c) $\max_{0 \le t \le 36} \beta(t)$ with $t > 9$ ;
d) No over-dispersion in the data: taking $\delta = 0.35 \max_t \beta(t)$, RIAs at distance larger than $\delta$ from the curve $\beta(t)$ were considered outliers and peptides with more than 25% outlier RIAs considered over-dispersed ;
e) $\lambda < 1$ ;
f) $\max_{0 \le t \le 36} \beta(t) < 0.2$ ;
g) Linear correlation between experimental data (RIAs) and $\beta(t)$ > 0.75 ;
h) At least two non-outlier RIAs available for $t \le 10$ ;
i) At least two non-outlier RIAs available for $t \ge 20$ ;
j) Build a loess piecewise polynomial approximation using all the non-outlier RIAs (R loess function), predict pointwise standard error (SE), count the proportion of times $t$ outside the loess ± 2SE, this proportion must be less than 25%.

Step 4: protein model

For a given protein, take all the observations that passed step 3. In case there are three or more, subject to R function boxplot.stats the $k_c$ parameters for all the observation individual models computed at step 3. Ignore the observations whose $k_c$ is deemed an outlier by boxplot.stats.

Then, since each observation might include a small vertical shift (see explanation above), we empirically aligned peptides before fitting the mathematical model on the pool of all the observations RIAs. We considered that the higher the heavy peptide signal the more reliable the observation. We thus first identified the most intense observation after its heavy MS intensity. One observation intensity was determined by taking the third quartile of the heavy MS signal between times 9h and 32h. Using the most intense observation as an anchor, we simply applied vertical shifts to all the other observations by minimizing weighted squared median distances between the anchor and the observation to align, the weight being proportional to the square root of the median of the heavy MS intensities of the observation to align.

Next, a first mathematical model was constructed as above with all the – pooled – observations RIAs. Subsequently, we determined outliers as above but with a slightly larger $\delta = 0.5 \max_t \beta(t)$ to acknowledge additional variability due to multiple observations. Observations comprised of >25% outliers were discarded from the protein model. They might be nonetheless correct as isoform, chain or active peptide with a different kinetics, meaning that they were maintained as correct as independent observation. Lastly, the mathematical model was recomputed with all the kept observations.

Step 5: bootstrap

In order to obtain an estimate of the variability in our observations (CI95), we applied a nonparametric balanced bootstrap using R library boot, 1,000 resamplings, and basic bootstrap intervals. Note that information obtained with this bootstrap step is not used in the multi-compartment modeling.

### Simplified model BUGS code

```
# declare ODE
solution[1:ngrid,1:ndim] <- ode(init[1:ndim],times[1:ngrid],
                                D(C[1:ndim],t),origin,tol)

# compartment 1: all organs through plasma
llambda_1 ~ dnorm(llambda_1.mu,llambda_1.prec)
lambda_1 <- exp(llambda_1)
lk_1 ~ dnorm(lk_1.mu,lk_1.prec)
k_1 <- exp(lk_1)
lk_10 ~ dnorm(lk_10.mu,lk_10.prec)
k_10 <- exp(lk_10)
lk_13 ~ dnorm(lk_13.mu,lk_13.prec)
k_13 <- exp(lk_13)

# compartment 2: CNS
llambda_2 ~ dnorm(llambda_2.mu,llambda_2.prec)
lambda_2 <- exp(llambda_2)
lk_2 ~ dnorm(lk_2.mu,lk_2.prec)
k_2 <- exp(lk_2)
lk_20 ~ dnorm(lk_20.mu,lk_20.prec)
k_20 <- exp(lk_20)
lk_23 ~ dnorm(lk_23.mu,lk_23.prec)
k_23 <- exp(lk_23)

# compartment 3: CSF
lk_31 ~ dnorm(lk_31.mu,lk_31.prec)
k_31 <- exp(lk_31)
lk_32 ~ dnorm(lk_32.mu,lk_32.prec)
k_32 <- exp(lk_32)

# ODE definition
ilc <- step(9-t)
D(C[1], t) <- (lambda_1*ilc - C[1]) * k_1
D(C[2], t) <- k_1*C[1] - (k_10+k_13)*C[2] + k_31*C[5]
D(C[3], t) <- (lambda_2*ilc - C[3]) * k_2
D(C[4], t) <- k_2*C[3] - (k_20+k_23)*C[4] + k_32*C[5]
D(C[5], t) <- k_13*C[2] + k_23*C[4] - (k_31+k_32)*C[5]

# residuals in plasma
for (i in offset[1]:(offset[2]-1)){
  j[i] <- i_times[i]
  mu.resid[i] <- solution[j[i],2]-shift[1]
  w.tau[i] <- tau[1]*weights[i]
  ratios[i] ~ dnorm(mu.resid[i],w.tau[i])
}
shift[1] ~ dnorm(shift.mu,shift.prec)
tau[1] ~ dgamma(0.001,0.001)

# residuals in CSF
for (i in offset[2]:(offset[3]-1)){
  j[i] <- i_times[i]
  mu.resid[i] <- solution[j[i],5]-shift[2]
  w.tau[i] <- tau[2]*weights[i]
  ratios[i] ~ dnorm(mu.resid[i],w.tau[i])
}
shift[2] ~ dnorm(shift.mu,shift.prec)
tau[2] ~ dgamma(0.001,0.001)
```

### Full model BUGS code

```
# declare ODE
solution[1:ngrid,1:ndim] <- ode(init[1:ndim],times[1:ngrid],
                                D(C[1:ndim],t),origin,tol)

# compartment 1: all organs through plasma
llambda_1 ~ dnorm(llambda_1.mu,llambda_1.prec)
lambda_1 <- exp(llambda_1)
lk_1 ~ dnorm(lk_1.mu,lk_1.prec)
k_1 <- exp(lk_1)
lk_10 ~ dnorm(lk_10.mu,lk_10.prec)
k_10 <- exp(lk_10)
lk_12 ~ dnorm(lk_12.mu,lk_12.prec)
k_12 <- exp(lk_12)
lk_13 ~ dnorm(lk_13.mu,lk_13.prec)
k_13 <- exp(lk_13)

# compartment 2: CNS
llambda_2 ~ dnorm(llambda_2.mu,llambda_2.prec)
lambda_2 <- exp(llambda_2)
lk_2 ~ dnorm(lk_2.mu,lk_2.prec)
k_2 <- exp(lk_2)
lk_20 ~ dnorm(lk_20.mu,lk_20.prec)
k_20 <- exp(lk_20)
lk_21 ~ dnorm(lk_21.mu,lk_21.prec)
k_21 <- exp(lk_21)
lk_23 ~ dnorm(lk_23.mu,lk_23.prec)
k_23 <- exp(lk_23)

# compartment 3: CSF
lk_30 ~ dnorm(lk_30.mu,lk_30.prec)
k_30 <- exp(lk_30)
lk_31 ~ dnorm(lk_31.mu,lk_31.prec)
k_31 <- exp(lk_31)
lk_32 ~ dnorm(lk_32.mu,lk_32.prec)
k_32 <- exp(lk_32)

# ODE definition
ilc <- step(9-t)
D(C[1], t) <- (lambda_1*ilc - C[1]) * k_1
D(C[2], t) <- k_1*C[1] - (k_10+k_13+k_12)*C[2] + k_31*C[5] + k_21*C[4]
D(C[3], t) <- (lambda_2*ilc - C[3]) * k_2
D(C[4], t) <- k_2*C[3] - (k_20+k_23+k_21)*C[4] + k_32*C[5] + k_12*C[2]
D(C[5], t) <- k_13*C[2] + k_23*C[4] - (k_31+k_32+k_30)*C[5]

# residuals in plasma
for (i in offset[1]:(offset[2]-1)){
  j[i] <- i_times[i]
  mu.resid[i] <- solution[j[i],2]-shift[1]
  w.tau[i] <- tau[1]*weights[i]
  ratios[i] ~ dnorm(mu.resid[i],w.tau[i])
}
shift[1] ~ dnorm(shift.mu,shift.prec)
tau[1] ~ dgamma(0.001,0.001)

# residuals in CSF
for (i in offset[2]:(offset[3]-1)){
  j[i] <- i_times[i]
  mu.resid[i] <- solution[j[i],5]-shift[2]
  w.tau[i] <- tau[2]*weights[i]
  ratios[i] ~ dnorm(mu.resid[i],w.tau[i])
}
shift[2] ~ dnorm(shift.mu,shift.prec)
tau[2] ~ dgamma(0.001,0.001)
```

### Supplementary Figures

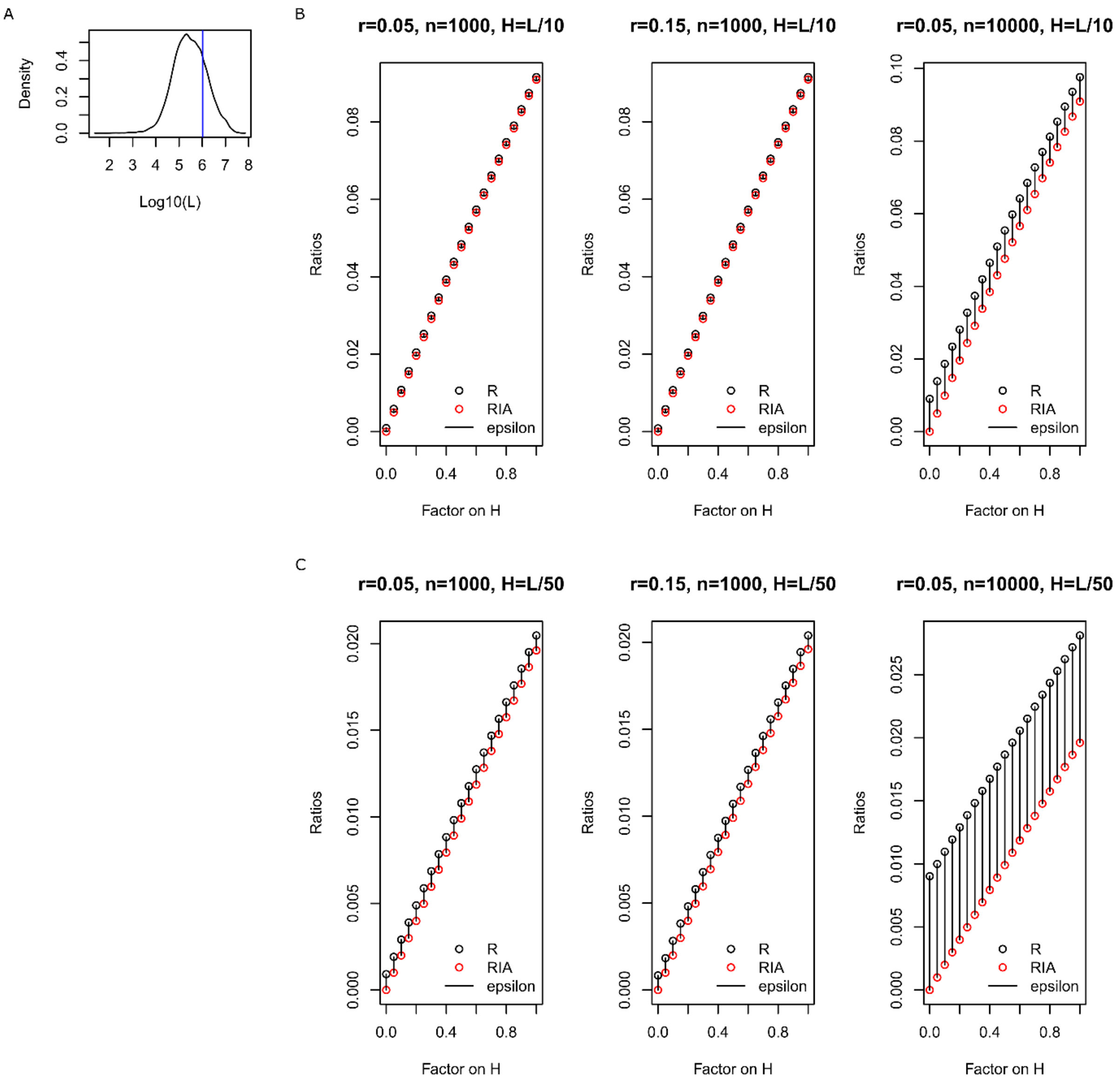


**Figure S1.** Simulations of the impact of multiplicative and additive noise on the observed ratios. (**A**) Distribution of the value of the light peptide signals $L$ observed for all the observations and time points, in all the unique peptides used for the protein models. We set $L = 1{,}036{,}522$ for the simulations, the mean value (vertical blue line). (**B**) Simulation of rather high ratios with a typical $H$ intensity equal to $L/10$. Different levels of additive and multiplicative noise were tested. We observe the expected vertical shifts at different magnitudes that primarily depend on the additive noise level, which was expected since we compute ratios. $R$ is the observed value, RIA is the exact value, and epsilon is $\varepsilon$ above, the approximation of the difference. To simulate different levels of tracer incorporation at different times, we multiplied the nominal $H$ by a factor between 0 and 1. We see that $\varepsilon$ is very accurate and almost constant as predicted. (**C**) Same simulations and observations for smaller RIAs with typical $H$ equal to $L/50$.

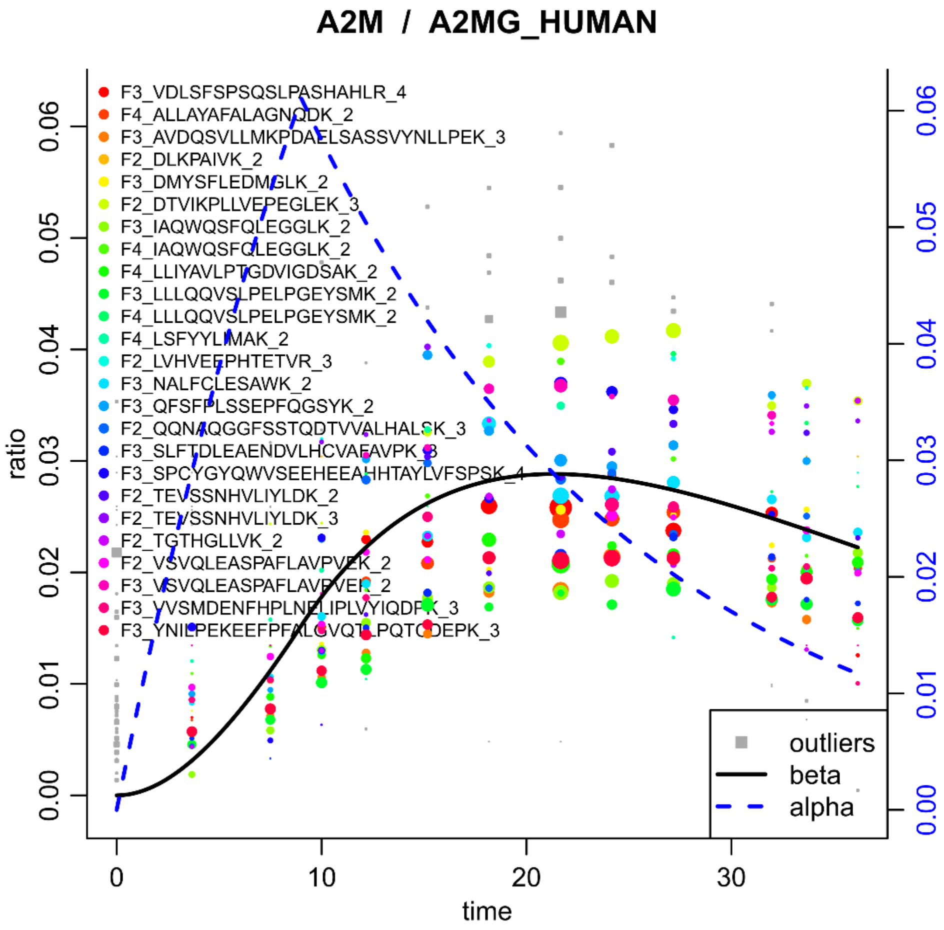


**Figure S2.** Example (Alpha-2-Macroglobulin, A2M) with uncorrected vertical shifts for each peptide.

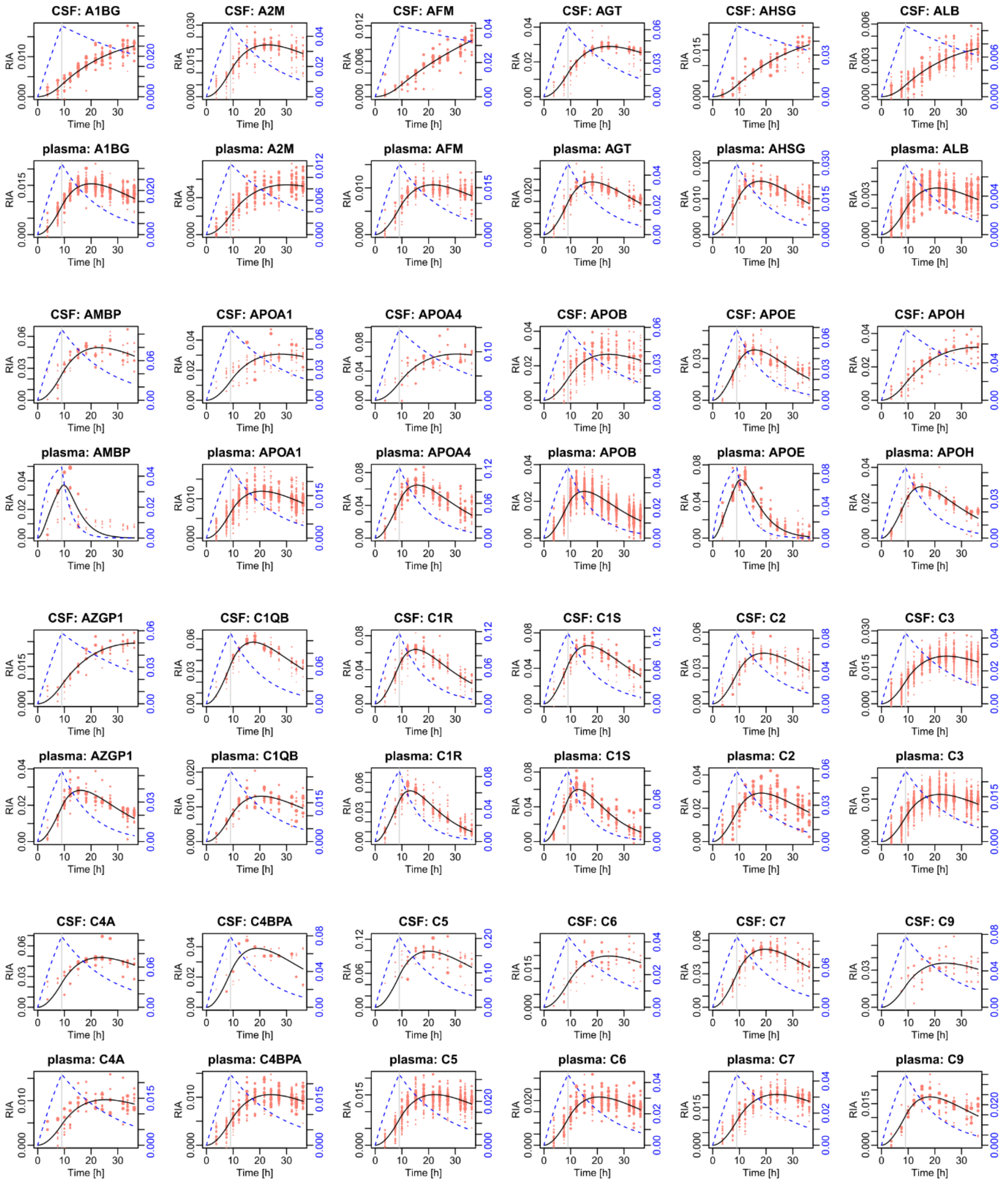


**Figure S3.** Protein dynamics (part 1) in plasma and CSF with independent models ($\beta(t)$ black solid curve, $\alpha(t)$ dashed blue curve).

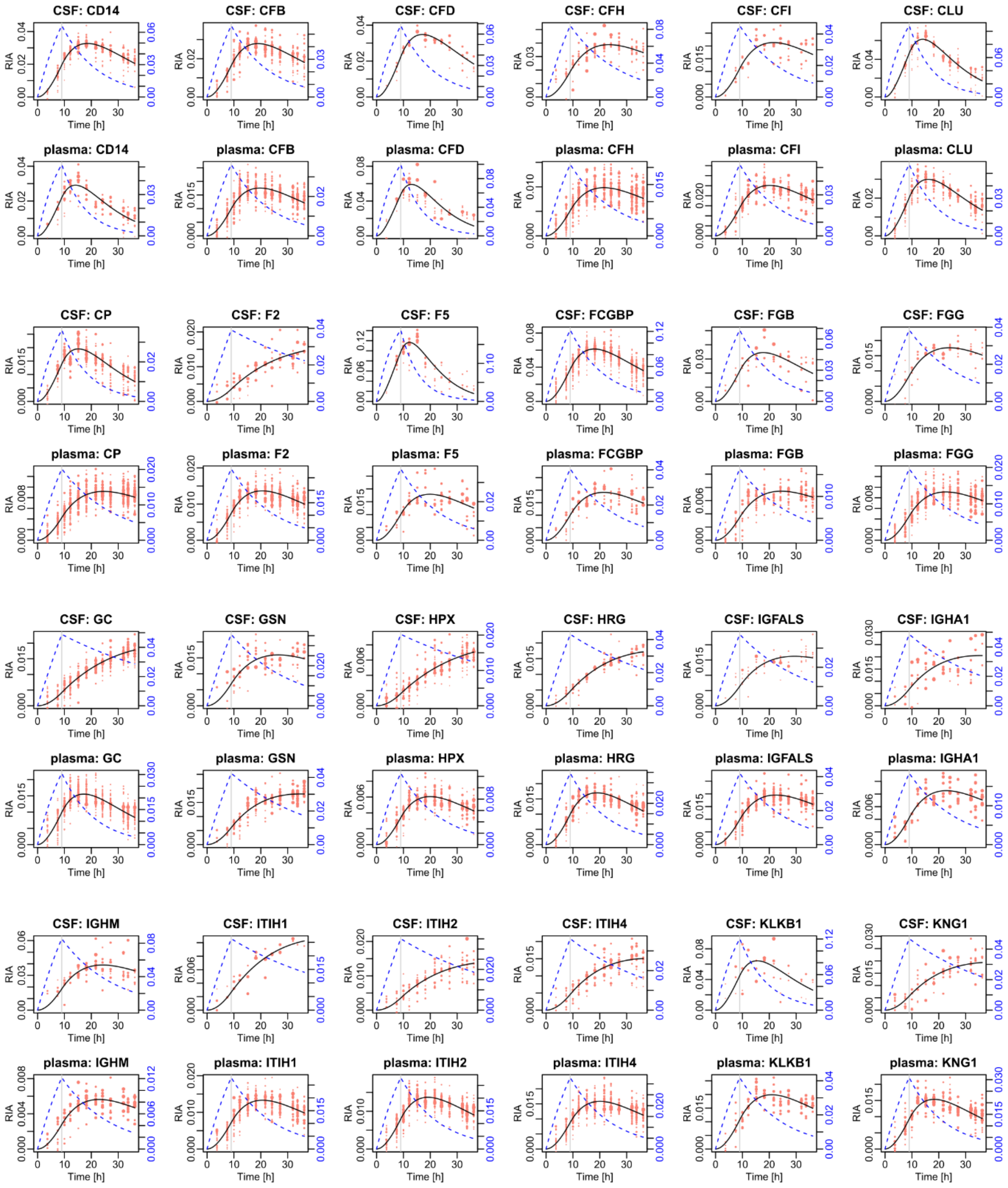


**Figure S4.** Protein dynamics (part 2) in plasma and CSF with independent models ($\beta(t)$ black solid curve, $\alpha(t)$ dashed blue curve).

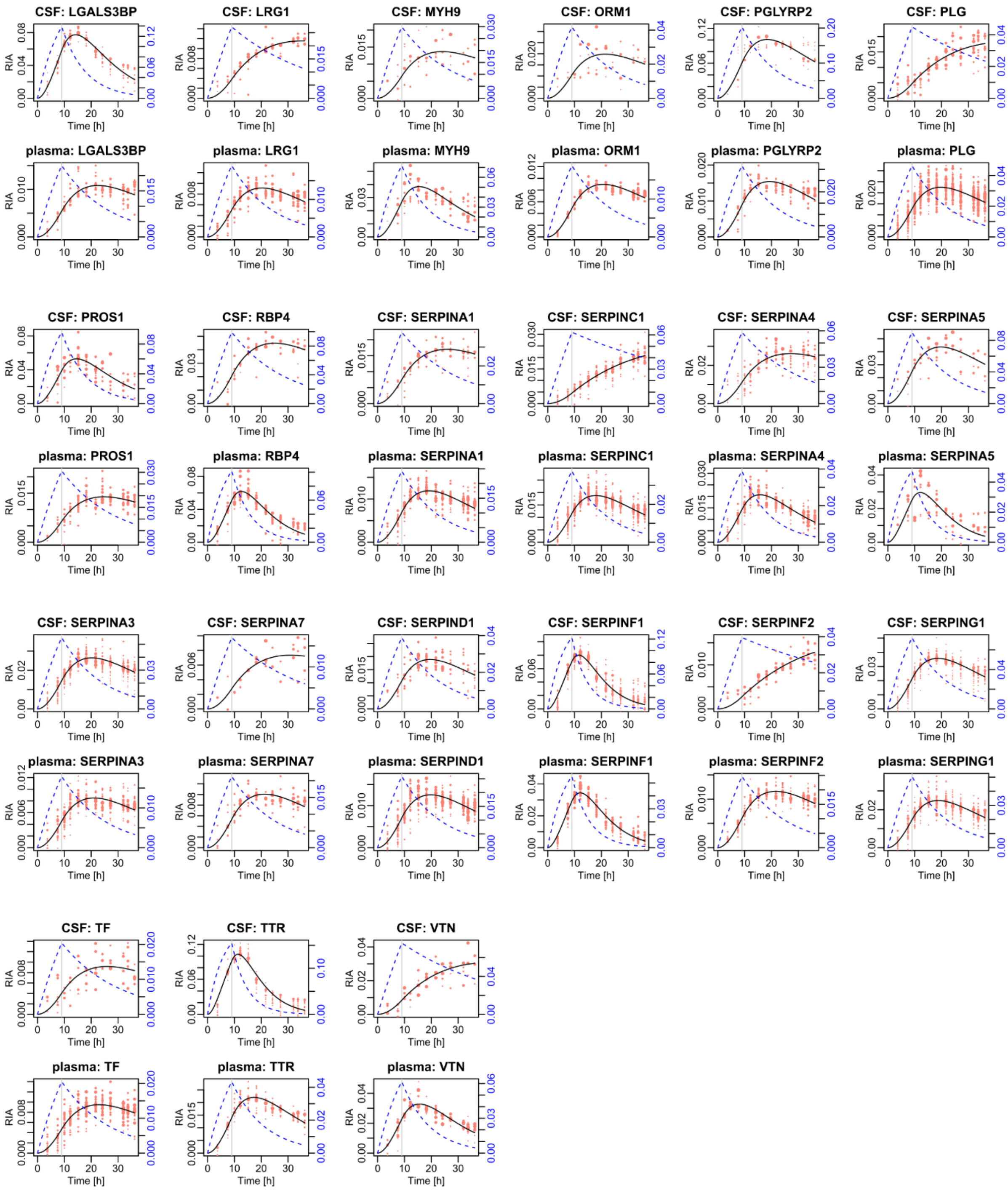


**Figure S5.** Protein dynamics (part 3) in plasma and CSF with independent models ($\beta(t)$ black solid curve, $\alpha(t)$ dashed blue curve).

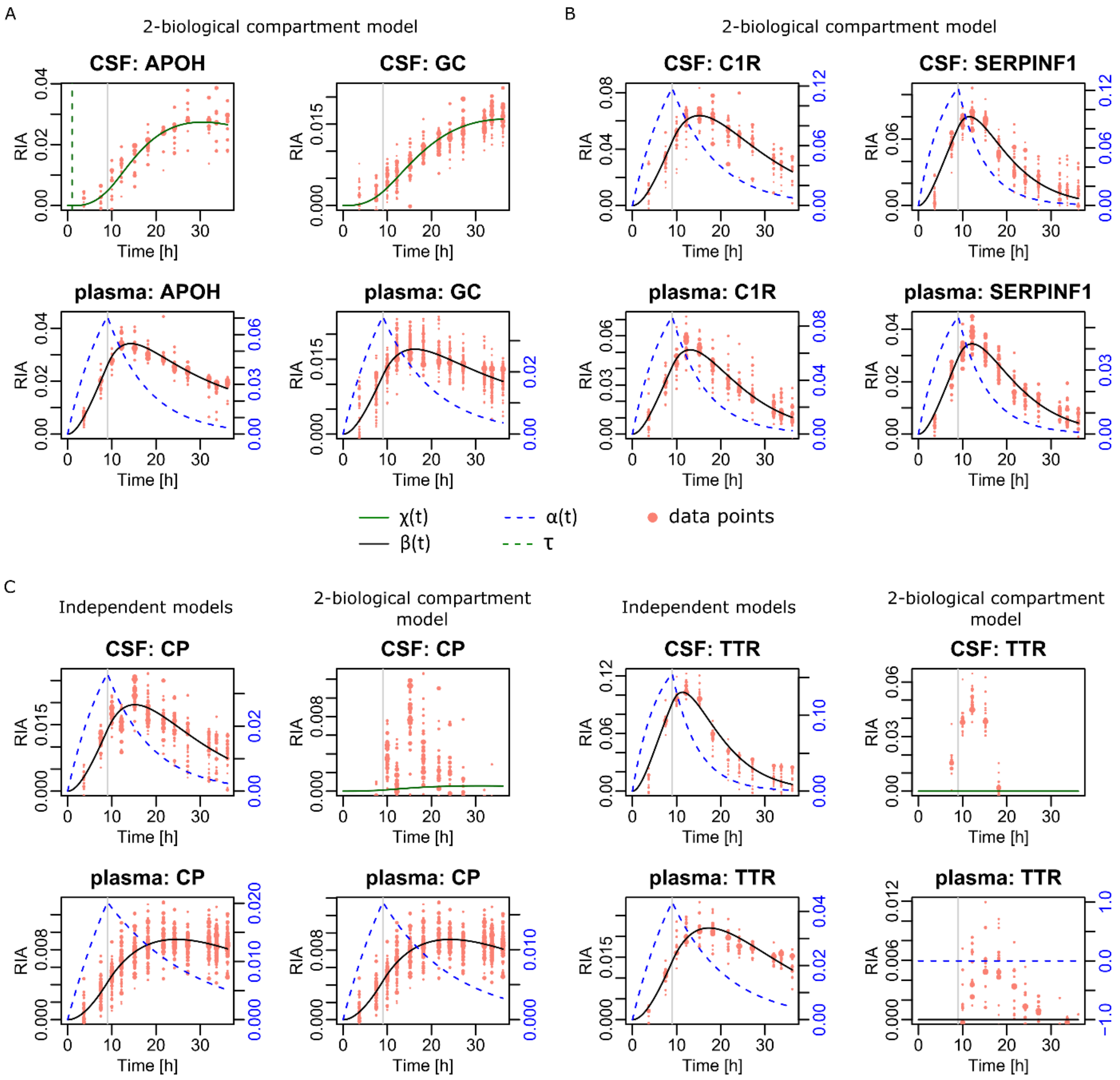


**Figure S6.** Additional examples with the simple, 2-biological compartment model. (**A**) Two successful cases with slower CSF dynamics. (**B**) Two successful cases with comparable dynamics. (**C**) Two failed cases with CSF faster dynamics. In each case, we show the independent fluid models on the left, and the 2-biological compartment model on the right.

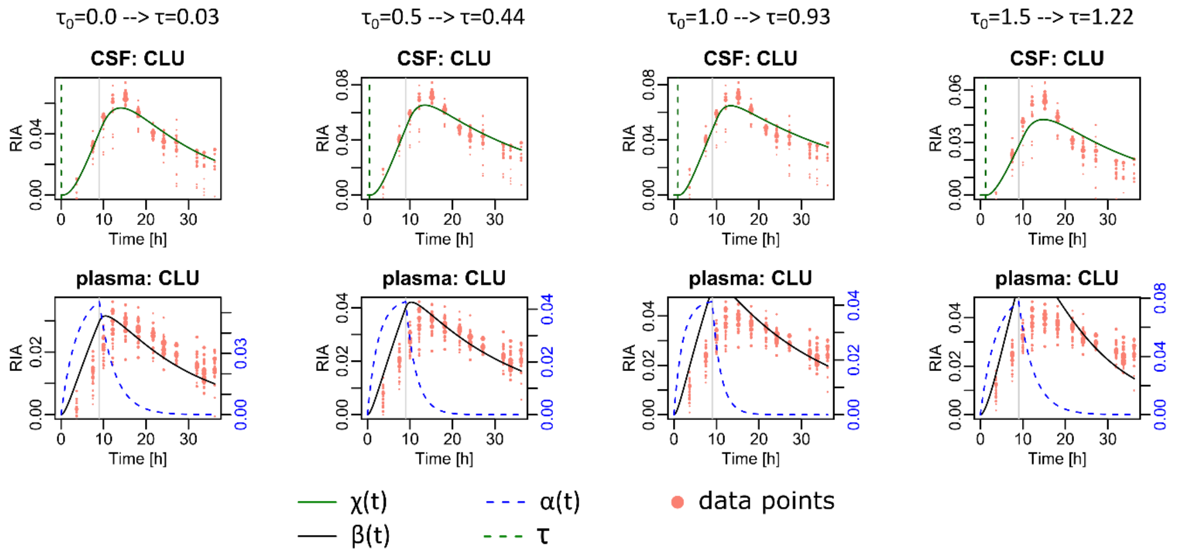


**Figure S7.** Results obtained with the delay differential model. Different initial values $\tau_0$ were tested for the delay, which yielded different delays $\tau$ after convergence. No improvement over the purely diffusive model with comparable accuracy for $\tau \leq 0.50$ hours, and worse for the other values. Larger initial values $\tau_0$ such as 3 or 9 hours yielded absurd solutions, much worse than values $\tau_0$ equal to 1 or 1.5 above, with $\chi(t)$ staying close to 0 over the whole time period (data not shown).

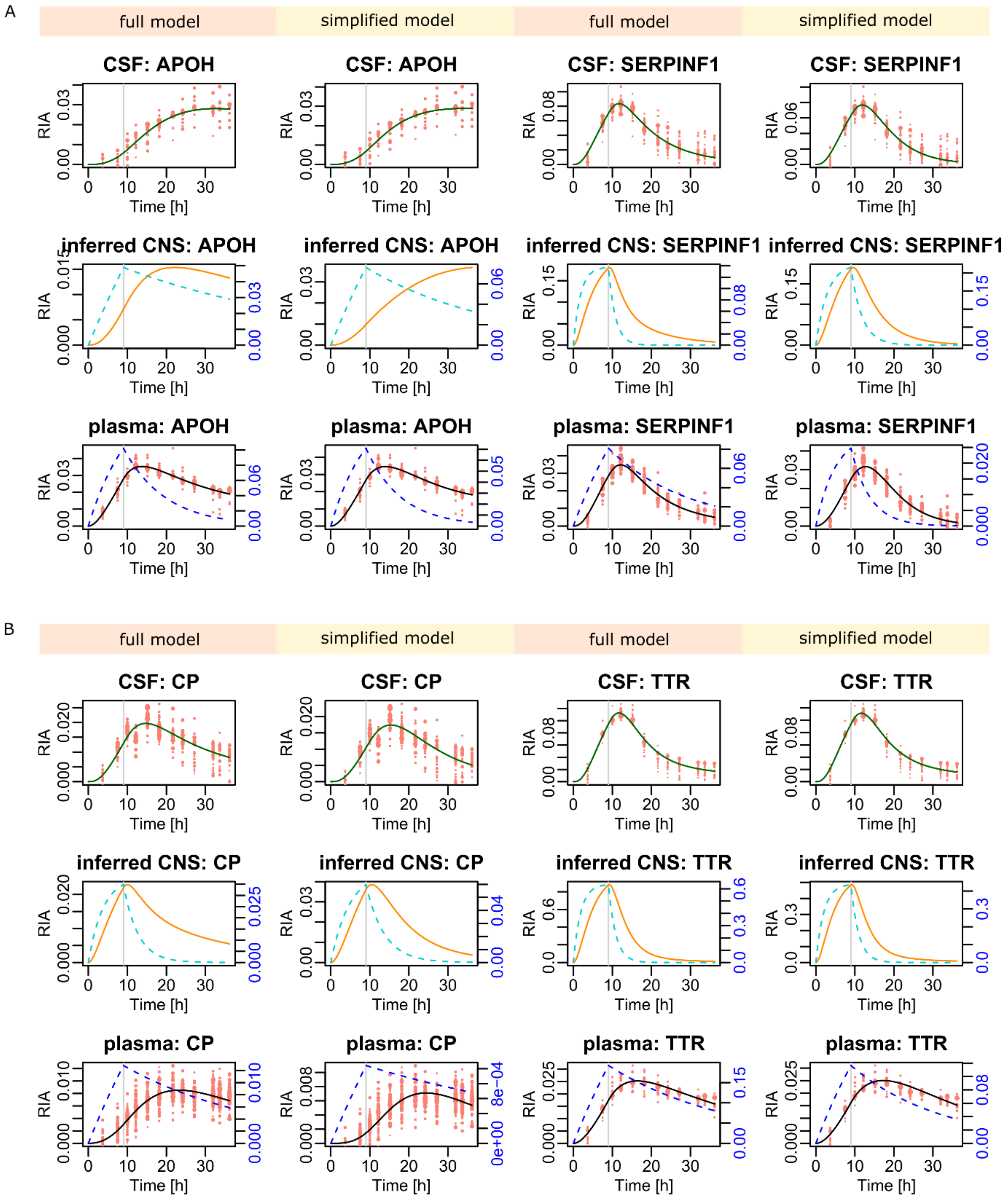


**Figure S8.** Additional examples with the 3- biological compartment models (full and simplified). (**A**) Two proteins with either slower CSF dynamics (Apolipoprotein H, APOH) or comparable dynamics (Serpin family F member 1, SERPINF1) for which the 2-biological compartment model achieved satisfying accuracy (Figure S4). (**B**) Two proteins (Ceruloplasmin, CP, and transthyretin, TTR) for which the 2-biological compartment model failed completely (Figure S4). We note the accurate fit achieved by both the full and the simplified models in panels (A) and (B) with rather similar solutions. The inferred CNS dynamics varies more between the full and simplified model solutions, an extreme case being APOH.
