## Supplementary figures and images for "Modeling the simultaneous dynamics of proteins in blood plasma and the cerebrospinal fluid in human *in vivo*"

### graph-model-plot.pdf

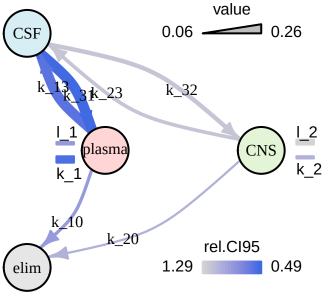

### graph-model-plot.pdf

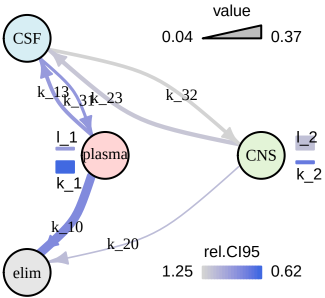

### graph-model-plot.pdf

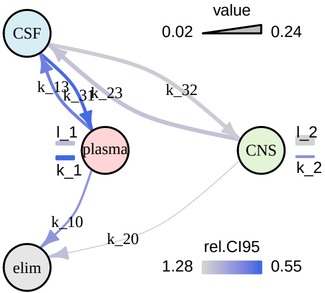

### graph-model-plot.pdf

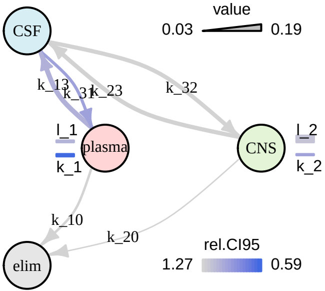

### graph-model-plot.pdf

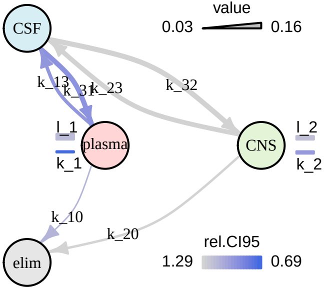

### graph-model-plot.pdf

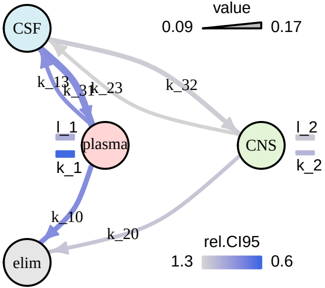

### graph-model-plot.pdf

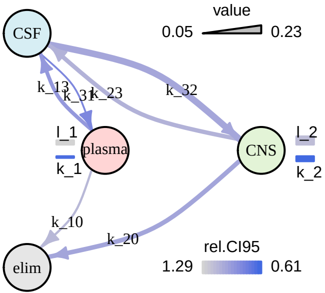

### graph-model-plot.pdf

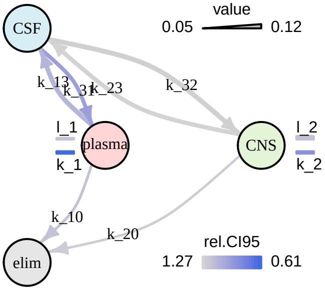

### graph-model-plot.pdf

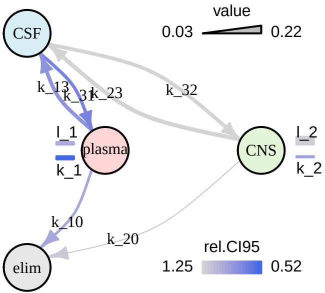

### graph-model-plot.pdf

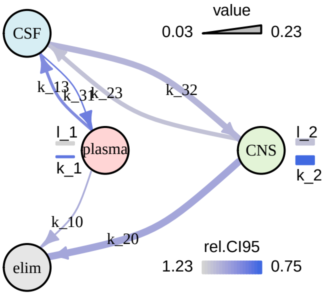

### graph-model-plot.pdf

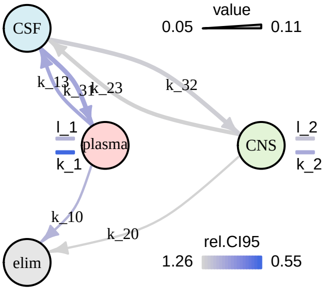

### graph-model-plot.pdf

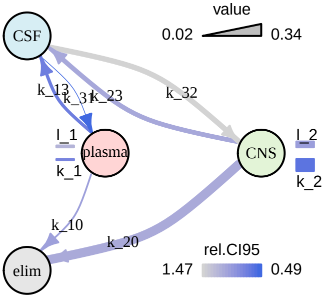

### graph-model-plot.pdf

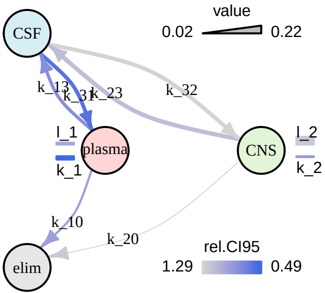

### graph-model-plot.pdf

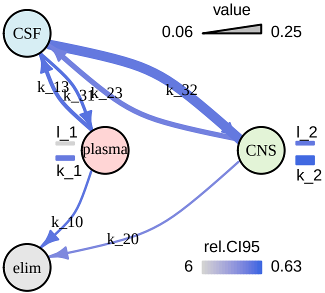

### graph-model-plot.pdf

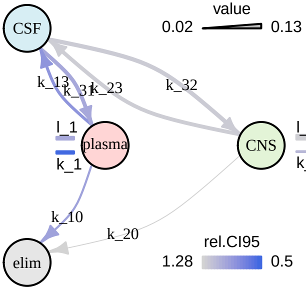

### graph-model-plot.pdf

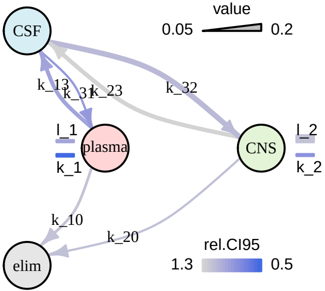

### graph-model-plot.pdf

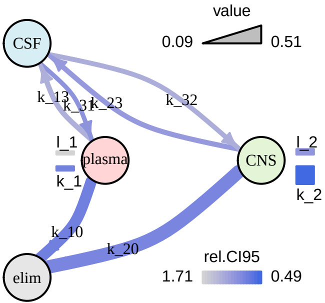

### graph-model-plot.pdf

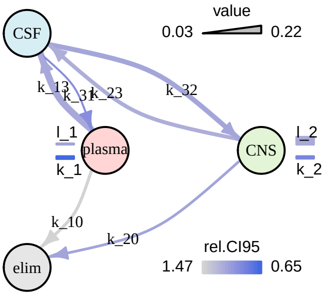

### graph-model-plot.pdf

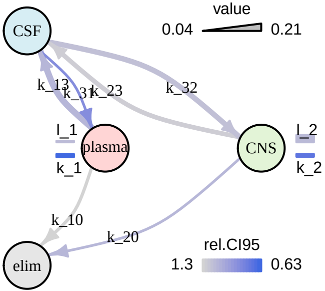

### graph-model-plot.pdf

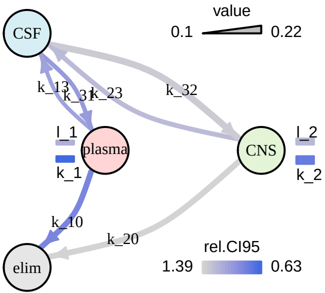

### graph-model-plot.pdf

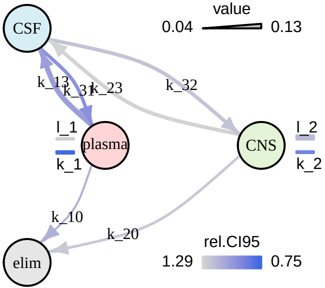

### graph-model-plot.pdf

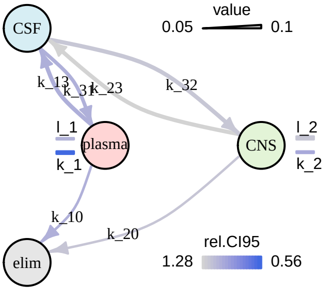

### graph-model-plot.pdf

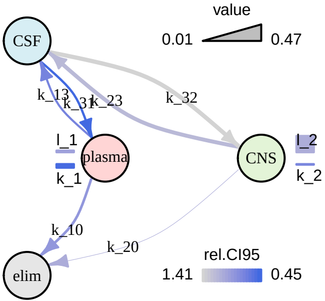

### graph-model-plot.pdf

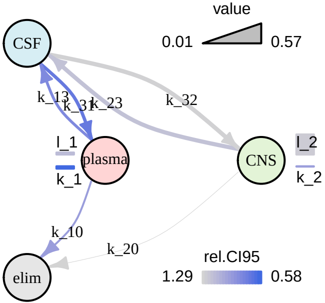

### graph-model-plot.pdf

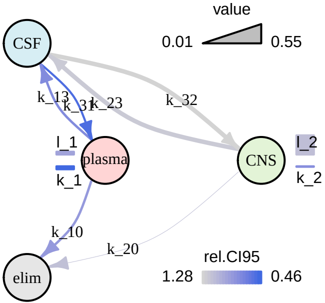

### graph-model-plot.pdf

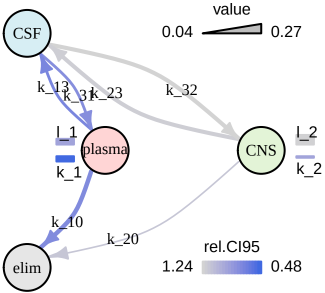

### graph-model-plot.pdf

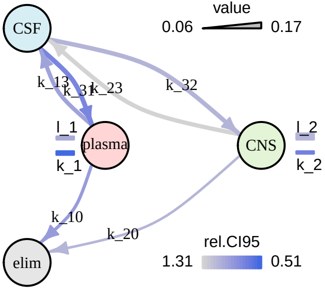

### graph-model-plot.pdf

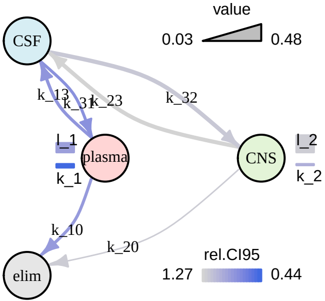

### graph-model-plot.pdf

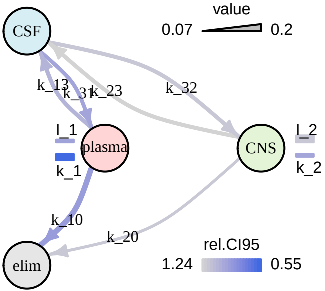

### graph-model-plot.pdf

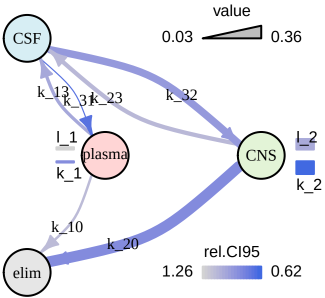

### graph-model-plot.pdf

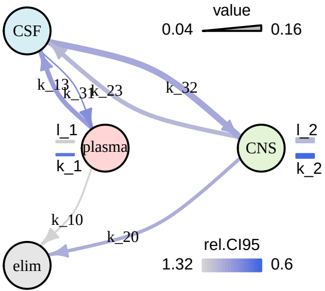

### graph-model-plot.pdf

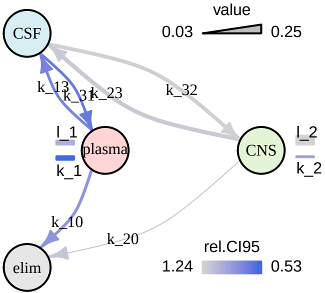

### graph-model-plot.pdf

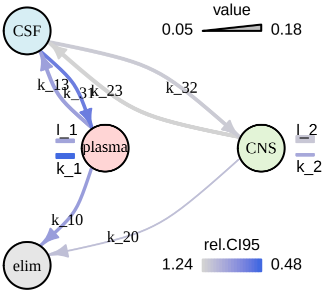

### model-plot.pdf

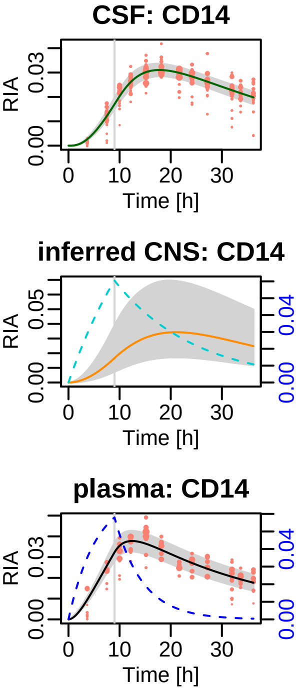

### model-plot.pdf

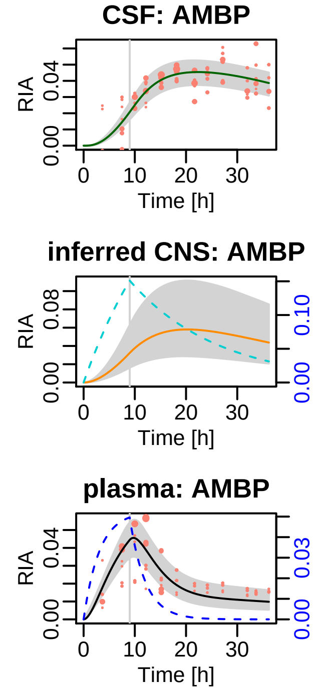

### model-plot.pdf

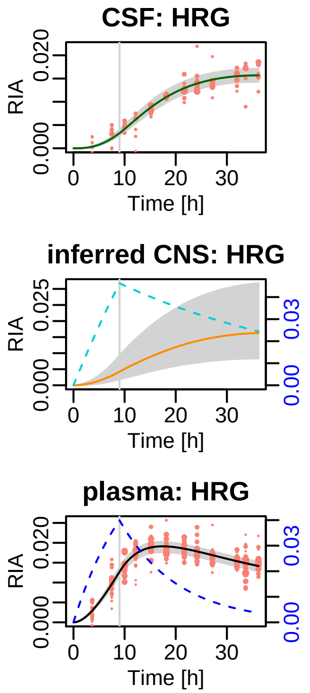

### model-plot.pdf

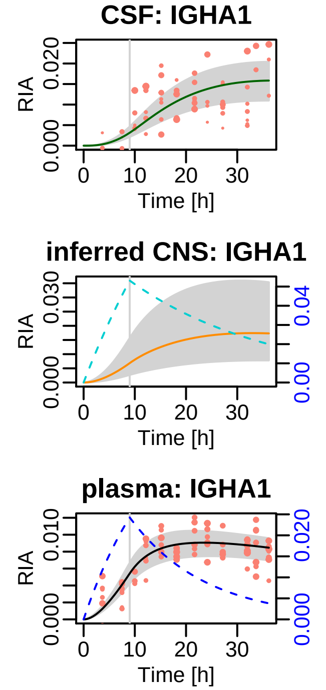

### model-plot.pdf

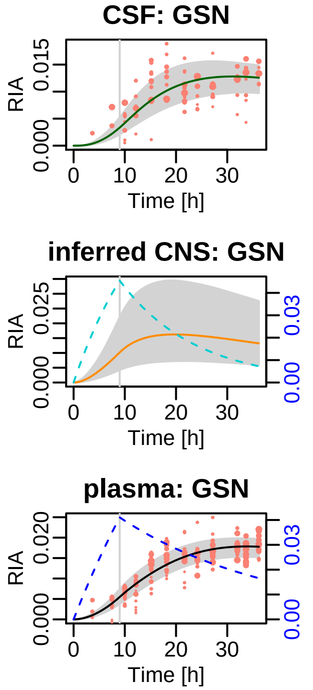

### model-plot.pdf

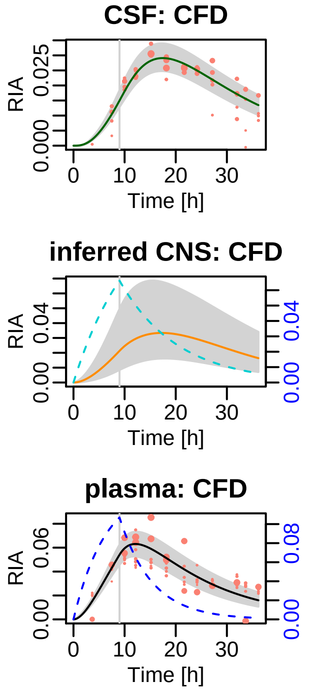

### model-plot.pdf

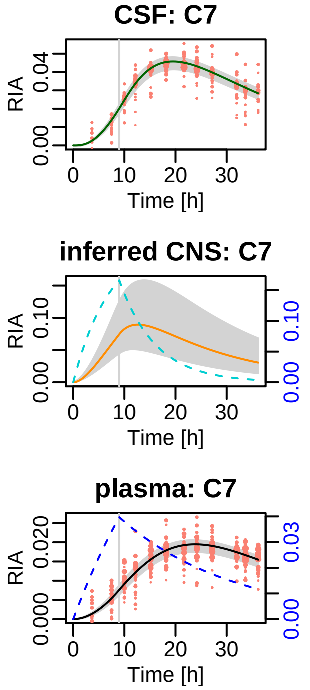

### model-plot.pdf

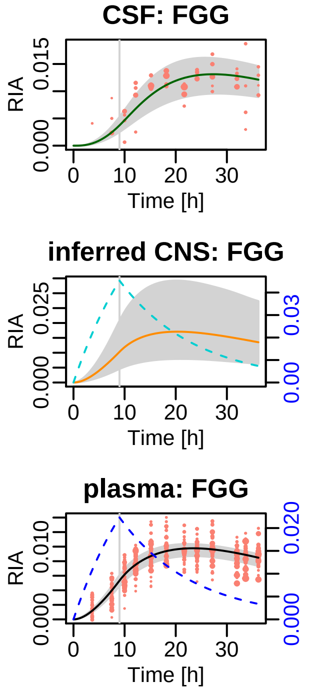

### model-plot.pdf

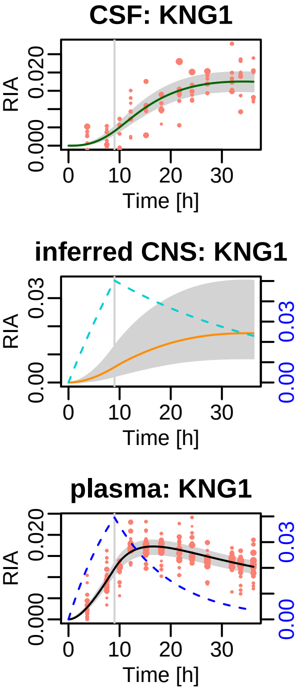

### model-plot.pdf

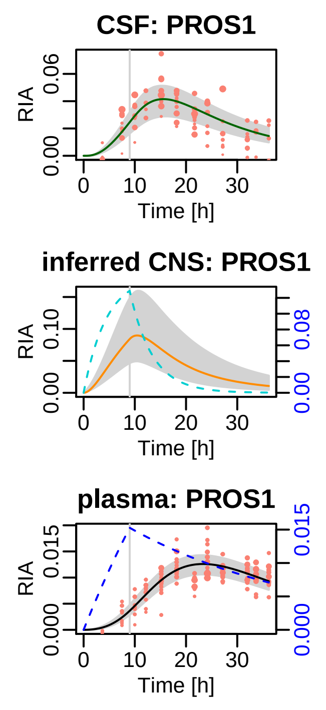

### model-plot.pdf

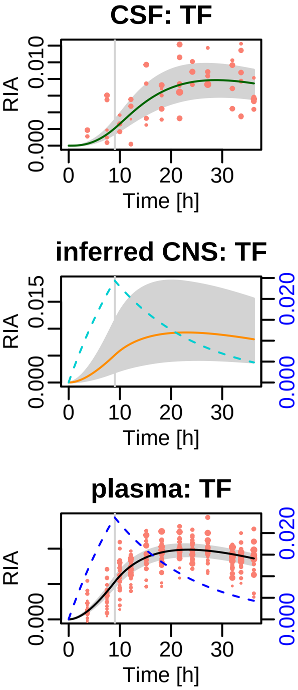

### model-plot.pdf

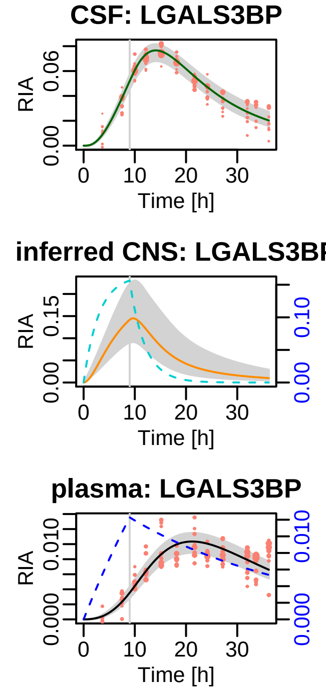

### model-plot.pdf

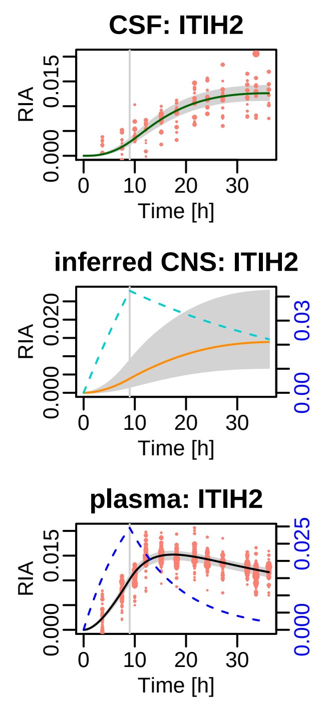

### model-plot.pdf

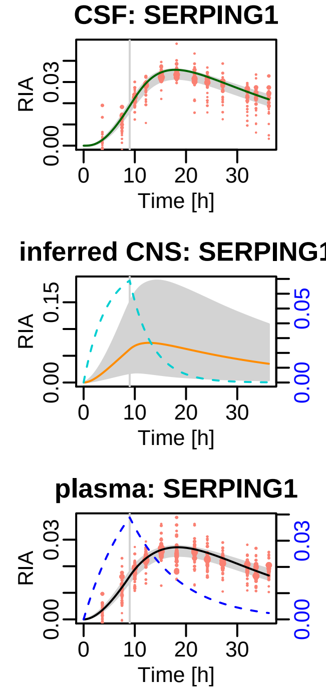

### model-plot.pdf

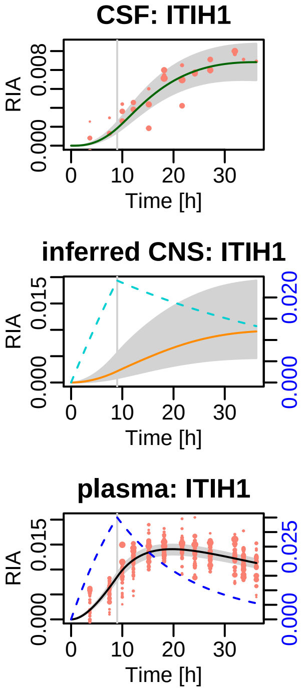

### model-plot.pdf

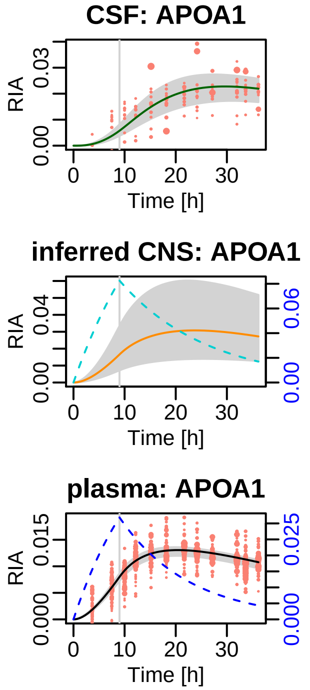

### model-plot.pdf

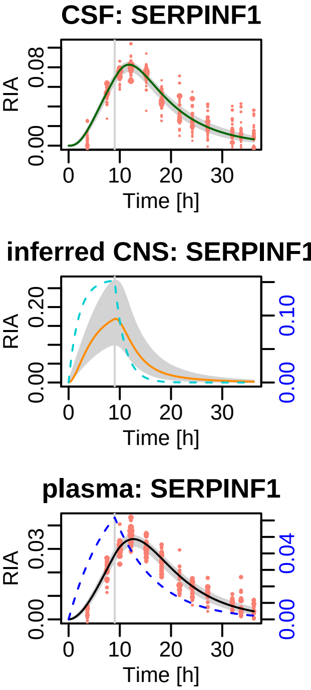

### model-plot.pdf

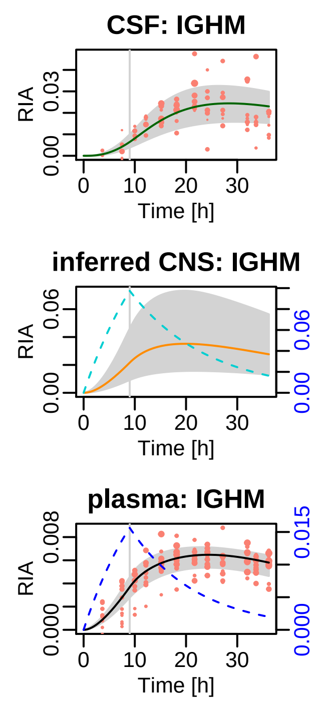

### model-plot.pdf

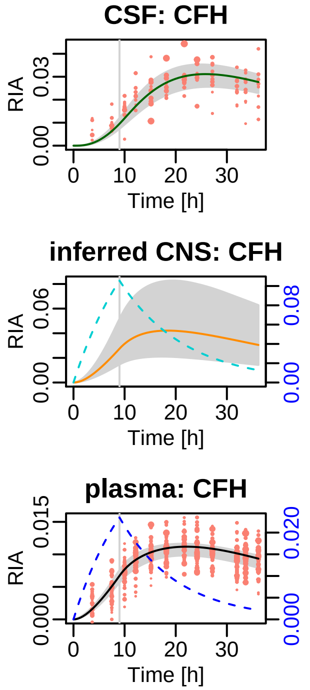

### model-plot.pdf

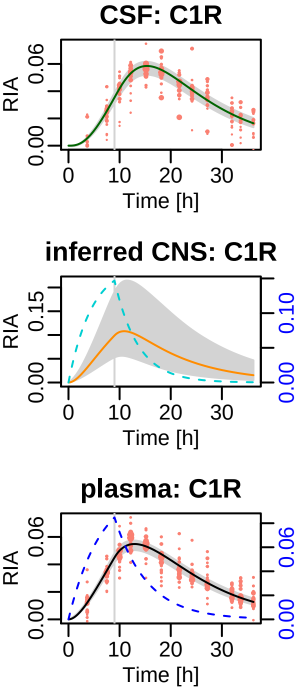

### model-plot.pdf

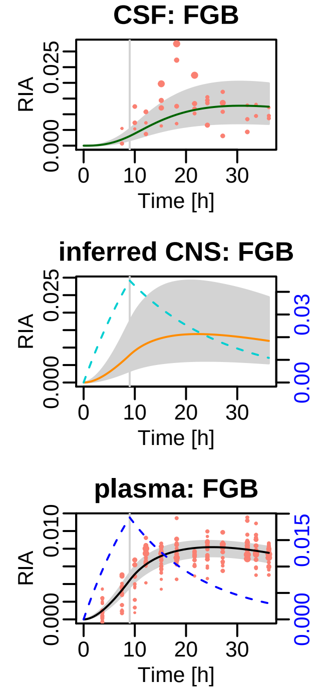

### model-plot.pdf

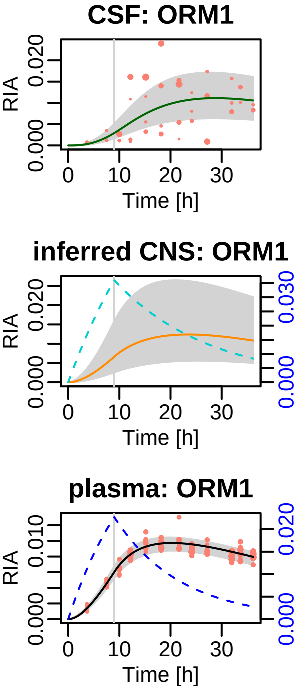

### model-plot.pdf

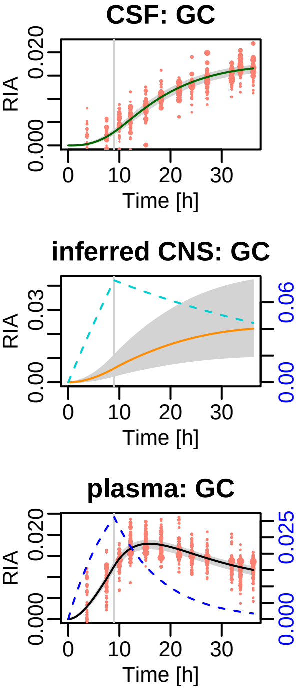

### model-plot.pdf

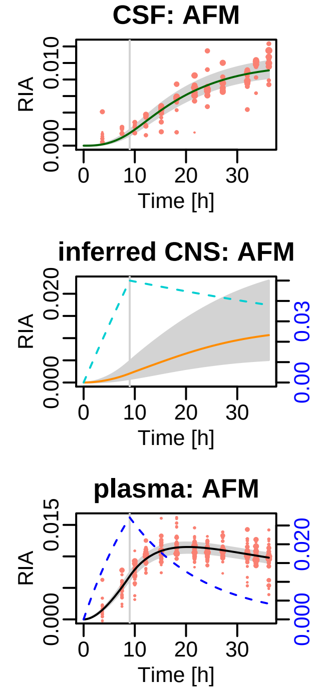

### model-plot.pdf

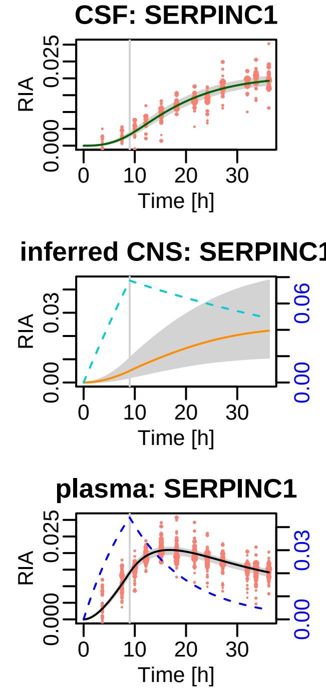

### model-plot.pdf

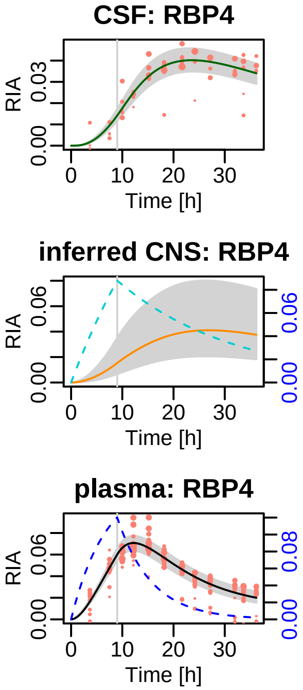

### model-plot.pdf

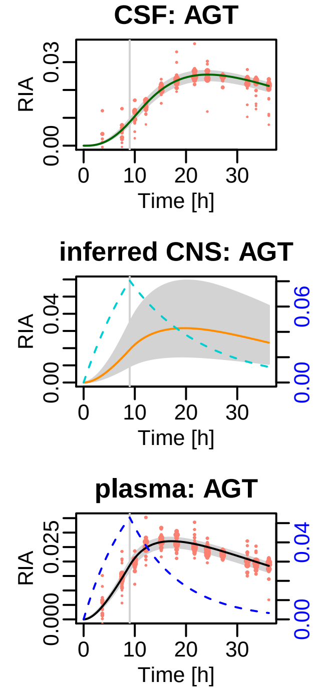

### model-plot.pdf

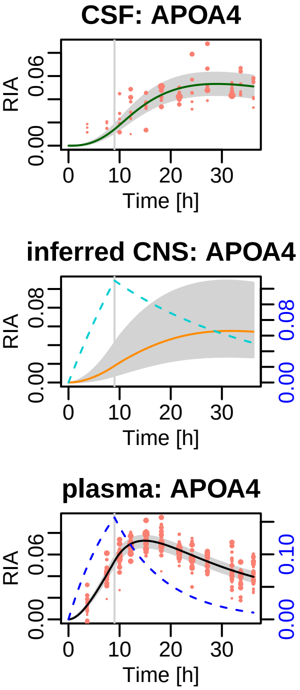

### model-plot.pdf

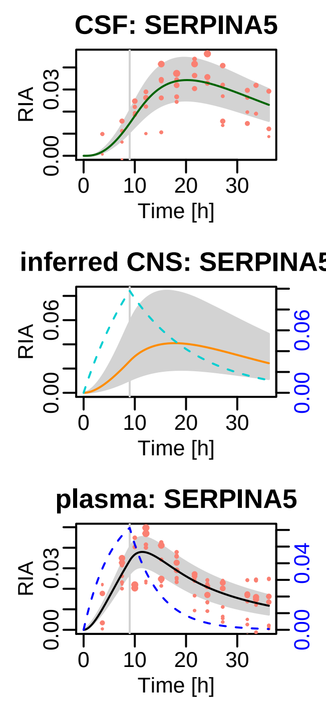

### model-plot.pdf

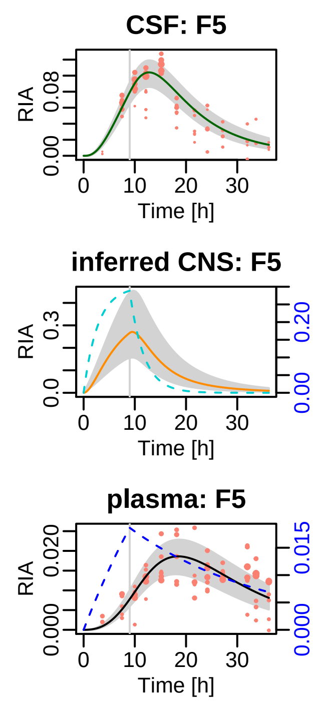

### model-plot.pdf

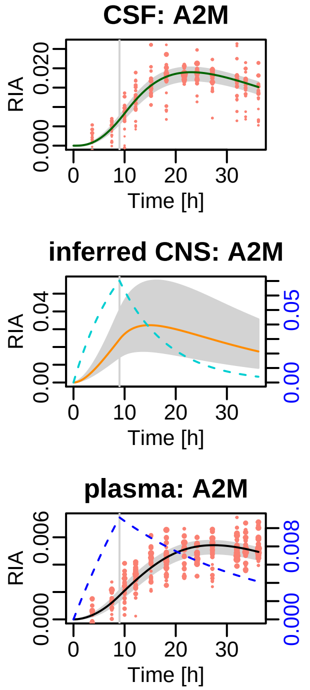

### model-plot.pdf

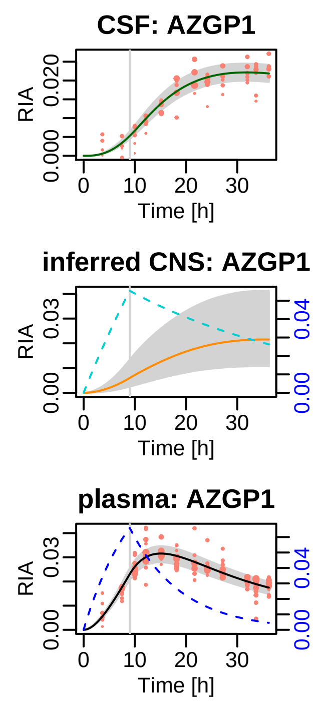

### model-plot.pdf

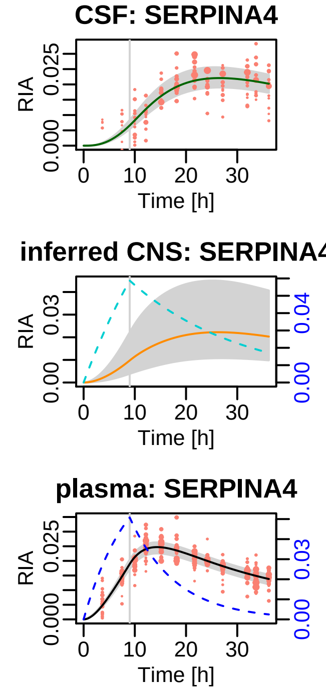

### model-plot.pdf

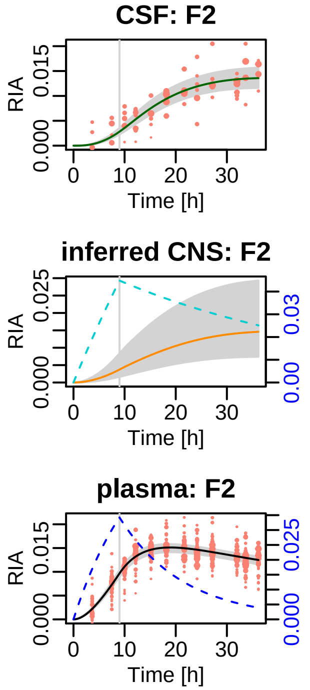
